# Enhancing the anti-tumor efficacy of Bispecific T cell engagers via cell surface glycocalyx editing

**DOI:** 10.1101/2022.05.22.492978

**Authors:** Zhuo Yang, Yingqin Hou, Geramie Grande, Chao Wang, Yujie Shi, Jaroslav Zak, Jong Hyun Cho, Dongfang Liu, John R. Teijaro, Richard A. Lerner, Peng Wu

## Abstract

Bispecific T-cell engager (BiTE)-based cancer therapies that activate the cytotoxic T cells of a patient’s own immune system have gained momentum with the recent FDA approval of Blinatumomab for treating B cell malignancies. However, this approach has had limited success in targeting solid tumors. Here, we report the development of BiTE-sialidase fusion proteins that enhance tumor cell susceptibility to BiTE-mediated cytolysis by T cells via targeted desialylation at the BiTE-induced T cell-tumor cell interface. Targeted desialylation results in better immunological synapse formation, T-cell activation and effector function. As a result, BiTE-sialidase fusion proteins show remarkably increased efficacy in inducing T-cell-dependent tumor cell cytolysis in response to target antigens compared to the parent BiTE molecules alone. This enhanced function is seen both *in vitro* and in *in vivo* xenograft and syngeneic solid tumor mouse models. Our findings highlight BiTE-sialidase fusion proteins as promising candidates for the development of next-generation bispecific T-cell engaging molecules for cancer immunotherapy.

## Introduction

A central theme in cancer immunotherapy is the activation of a patient’s own immune system for tumor control. Bispecific T cell engagers (BiTEs) are off-the-shelf agents that recruit endogenous CD8^+^ and CD4^+^ T cells capable of eradicating tumor cells in a manner that is independent of the major histocompatibility complex (MHC)^1^. A BiTE molecule consists of two single-chain variable fragments (scFvs), one targeting a tumor-associated antigen, while the other binds to CD3 on T cells. These two scFvs are covalently connected by a small linker peptide. Blinatumomab, which targets the CD19 antigen present on B cells, is the first BiTE approved by the US Food and Drug Administration (FDA) and is used to treat B-cell precursor acute lymphoblastic leukemia (ALL) in patients with residual cancer following chemotherapy^2^.

However, as with most T cell-based therapies, the promise of BiTEs in the treatment of solid tumors has yet to be realized^3^. In addition to the problem of limited penetration into the tumor tissue, T cell-based therapies must overcome the immunosuppressive tumor microenvironment, where T-cell suppression is orchestrated by tumor cells and the neighboring stromal myeloid and lymphoid cells^4^. In this unique microenvironment, limited availability of nutrients and accumulated metabolic waste products lead to alterations in cell-surface epitopes of both tumor and immune cells, which subsequently alters their interactions and ultimately leads to T cell exhaustion and poor tumor control^5^. Therefore, enabling approaches that target the molecular and cellular components of the immunosuppressive tumor microenvironment may transform T cell-based cancer treatments, including those based on T cell-engaging technologies.

Aberrant glycosylation is a hallmark of cancer^6–9^. Tumor cells often upregulate characteristic glycoforms with terminal sialic acid (hypersialylation), which can then act as glyco-immune checkpoints to suppress immune activation^10,11^. Sialosides attenuate immune cell activation and effector function by recruiting sialic acid-binding Ig-like lectins (Siglecs) that are found on most leukocytes to the immunological synapse, where they can trigger inhibitory signaling^12,13^. In addition, sialo-glycans expressed on T cells and antigen-presenting cells (APCs) may interact with CD28 on T cells to compete with its binding to CD80 on APCs, resulting in suppression of the co-stimulation required for T cell activation and survival^14^.

Previous studies by Adema and coworkers demonstrated that selective inhibition of cell-surface sialylation in the tumor microenvironment via intra-tumoral administration of an unnatural sialic acid mimic, P-3Fax-Neu5Ac, that inhibits sialyltransferases is a powerful intervention that potentiates cancer killing by T cells, while reducing the infiltrating regulatory T cells and myeloid suppressor cells^15^. Sialoside blockade enhanced antigen-specific CD8+ T cell-mediated cytolysis of tumor cells, in part by facilitating clustering of tumor cells with T cells. Inspired by this and other work indicating the importance of hyper-sialylation in suppressing T cell-induced killing^16^, herein, we report the development of BiTE-sialidase fusion proteins that can remove sialo-glycans at the T cell-tumor cell interface engaged by BiTE moledules, leading to the increase of T cell-dependent tumor cell cytolysis. We demonstrate that the enhanced tumor cell cytolysis is independent of the inhibitory sialo-glycan-Siglec signaling but results from a stronger immunological synapse formation induced by BiTEs. Furthermore, in multiple preclinical models of liquid and solid tumors, we show that BiTE-sialidase fusion proteins exhibit superior efficacy in the *in vivo* control of tumor proliferation and extension of survival in comparison with the parent BiTE molecules.

## Results

### Removal of sialic acids on the surface of tumor cells enhances BiTE-mediated tumor cell killing by T cells

To evaluate whether desialylation may enhance the susceptibility of tumor cells to BiTE-mediated cytotoxicity by T cells, we first constructed a BiTE molecule from a high-affinity HER2-targeting scFv 4D5 derived from Herceptin (K_D_=5nM) and a moderate-affinity human CD3-targeting scFv (K_D_=120nM, the clone TR66 based on which blinatumomab was constructed) (ref) (4D5 BiTE). We then treated HER2 positive SK-BR-3 human breast cancer cells with a sialidase derived from *Bifidobacterium longum subspecies infantis (B. infantis)* to remove cell-surface sialic acids. Expressed by human commensal bacteria *B. infantis*, this sialidase is known to hydrolyze both α2-3 and α2-6-linked sialosides with high efficiency.

Staining with FITC-*Sambucus nigra agglutinin* (SNA), that binds preferentially to sialic acid attached to terminal galactose in an α-2,6 linkage, confirmed the success of cell-surface desialylation (Figure 1a). Next, we incubated SK-BR-3 cells with 4D5 BiTE and PBMCs from healthy human donors in the presence or absence of *B. infantis* sialidase. At different effector (E) to target (T) ratios the addition of sialidase markedly potentiated the 4D5 BiTE-induced tumor cell killing by T cells in comparison with 4D5 BiTEs alone (Figure 1b and Figure S1a). The improvement of the BiTE-induced killing was sialidase dose-dependent (Figure S1b). A similar trend was observed when another HER2 expressing breast cancer cell line, MCF-7, was used as the target cell and when 4D5 BiTE treatment was combined with a sialylation inhibitor, P-3Fax-Neu5Ac (Figure 1b and 1c). To verify the enhancement of BiTE-induced killing following sialidase treatment, another BiTE molecule, PSMA BiTE, that targets Prostate-Specific Membrane Antigen (PSMA) was constructed. Again, strong cytotoxicity enhancement was observed when sialidase and BiTEs were added simultaneously (Figure 1d). To confirm that the effect of BiTEs on tumor cell killing is mediated through their interaction with T cells, we repeated the killing assay with MCF-7 cells using purified T cells. To our satisfaction, the sialidase treatment indeed led to better BiTE-mediated target cell killing (Figure S1c). Moreover, the addition of the sialidase also significantly enhanced IFN-γ secretion and T cell activation (Figure 1e and f).

**Figure 1:**
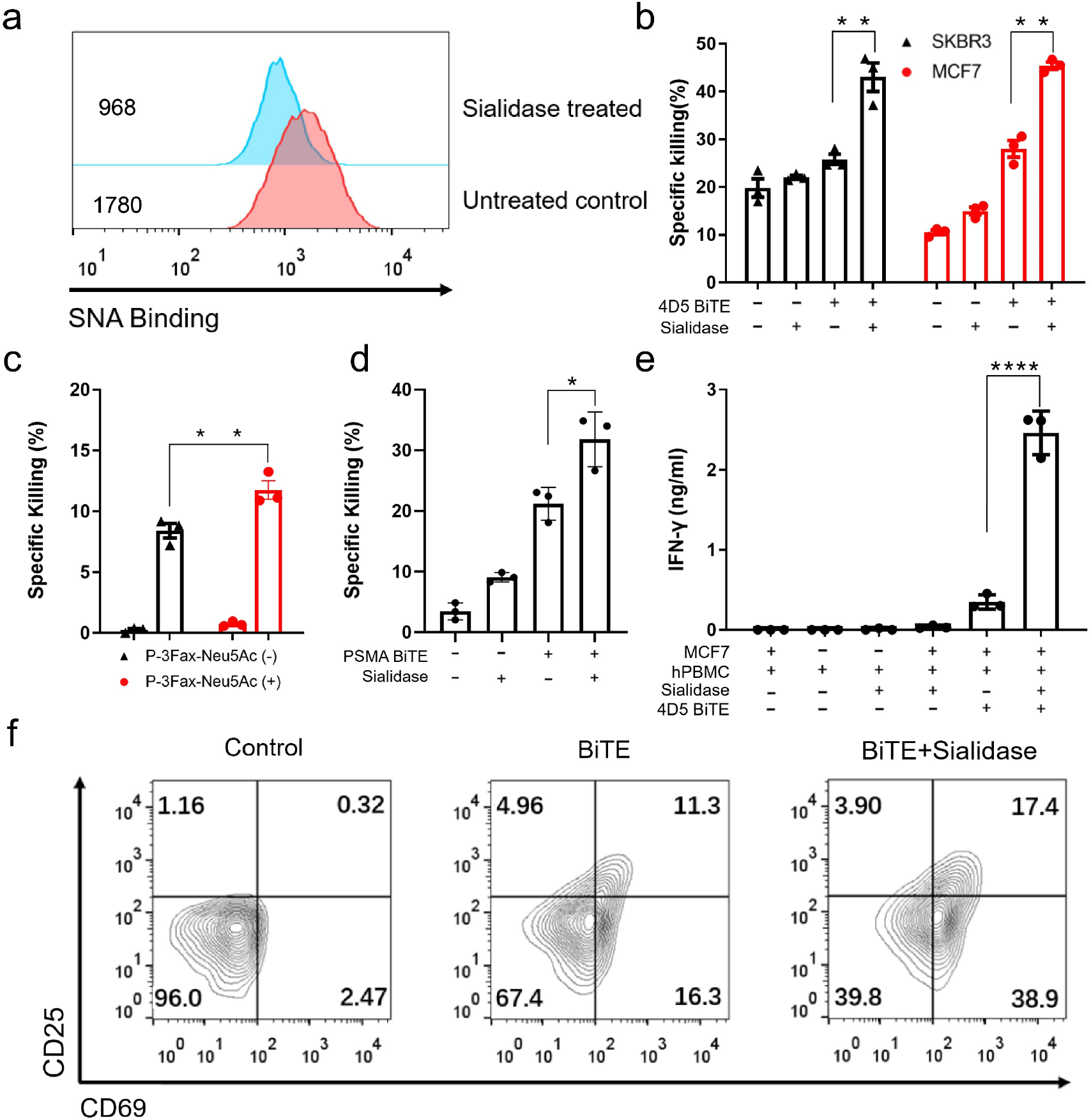
Sialic acid removal enhances BiTE-induced T cell cytotoxicity and activation (a) Measuring sialic acid levels on the surface of SK-BR-3 cells after the treatment of *B. infantis* sialidase with sialic acid binding lectin SNA. Mean fluorescent intensity (MFI) is displayed on the figure. (b) Killing of SK-BR-3 cells and MCF7 cells induced by 4D5 BiTEs with or without the sialidase treatment. (c) Killing of SK-BR-3 cells induced by 4D5 BiTEs + hPBMCs with or without prior treatment with 100 μM sialylation inhibitor P-3FAX-Neu5Ac. (d) Killing of PSMA positive PC3 cells induced by PSMA targeting BiTEs with or without the sialidase treatment. (e) IFN-γ release was measured as an indicator of BiTE-induced T cell activation by incubating MCF7 cells with 4D5 BiTEs + hPBMCs with or without the addition of sialidase. (f) CD25 and CD69 expression level was measured in T cells with or without the presence of 4D5 BiTEs and sialidase. The E to T ratio used for all experiments in Figure 1 is 5 to 1. Sialidase was added at concentration of 15 μg/ml. Mean values show three independent experiments with standard error of the mean (SEM) as error bars. For statistical analysis, unpaired Student t test with Welch correction was applied (\**P* < 0.05, \*\**P* < 0.01, \*\*\**P* < 0.001 and \*\*\*\**P* < 0.0001).

### Tumor cell desialylation promotes stronger BiTE-mediated immune synapse (IS) formation between T cells and tumor cells

Through their interaction with sialylated glycans aberrantly expressed on tumor cells, immune cell-associated Siglecs (sialic acid-binding immunoglobulin-type lectin) trigger signaling cascades to suppress immune cell activation and effector function^17,18^. Therefore, to probe the mechanism underlying the potentiation of BiTE-induced cytotoxicity by desialylation, we first investigated if the sialoglycan-Siglec inhibitory pathway is involved. Previously, Stanczak et al. observed increased T cell-dependent cytotoxicity of blinatumomab, a CD19-targeting BiTE, and catumaxomab, an Epithelial Cell Adhesion Molecule (EpCAM) targeting trifunctional antibody, against target tumor cells deficient in sialic acid^16^. They argued that the increased potency is due to the alleviation of the inhibitory signals mediated by the sialoglycan-Siglec-9 interaction. However, consistent with previous reports, we found that T cells from PMBCs of healthy donors expressed negligible levels of Siglec-7 and Siglec-9 as compared to their CD3 negative counterparts that mainly consist of B cells, NK cells, monocytes and dendritic cells (Figure 2b). Nevertheless, we did observe a slight up-regulation of both Siglec-9 and Siglec-7 following T cell activation. When compared to the expression of these Siglecs on freshly isolated CD3 negative cells, the expression of Siglec-7 and −9 on activated T cells was still minimal (Figure 2a). To further test if the Siglec-9 inhibitory pathway played a role in BiTE-induced T cell killing, a Siglec-9 blocking antibody was added with 4D5 BiTEs and the level of target cell killing was analyzed. In contrast to sialidase addition, blocking of the Siglec-9 signals did not increase cytotoxicity, indicating a negligible role of Siglec-9 in the BiTE-induced T cell killing (Figure 2c). Although the killing process mediated by BiTEs is often considered independent of the CD28 costimulatory signal, several studies have reported therapeutic advantages when T cell engager molecules are coupled with an additional CD28 recruiting moiety^19–21^. In addition, as demonstrated by Paulson and coworkers, sialoglycans can block the interaction of CD28 with CD80, resulting in dampened costimulatory signaling^14^. To investigate whether the enhanced BiTE-induced killing seen with desialylation is affected by CD28 co-stimulation, the CD28-CD80 interaction was blocked by adding a high affinity ligand of CD80, recombinant human CTLA-4. However, even with the addition of high concentrations of CTLA-4, no change in the enhanced cytolysis was detected (Figure 2d).

**Figure 2:**
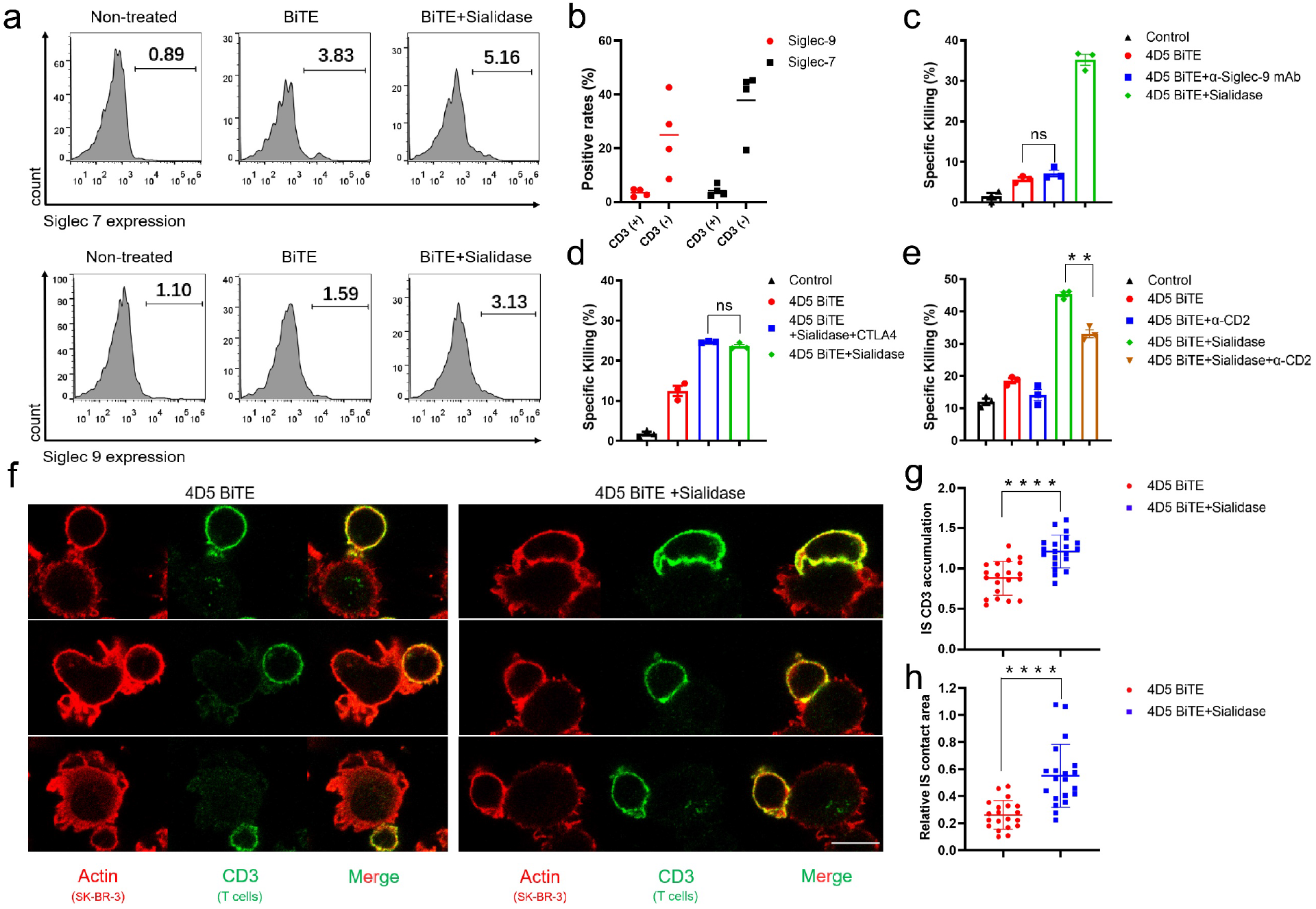
Desialylation promotes stronger BiTE-mediated immune synapse (IS) formation rather than suppressing the inhibitory Siglec signaling. (a) Siglec-7 and −9 expression levels were measured on human T cells with or without BiTE-induced activation, and with or without sialidase treatment. (b) Siglec-7 and −9 expression levels were measured on T cells and CD3 negative cells in PBMCs from healthy human donors. (c) SK-BR-3 cell killing induced by 4D5 BiTEs was measured with or without the addition of sialidase or anti-siglec-9 antibody. (d and e) SK-BR-3 cell killing induced by 4D5 BiTEs was measured in the presence or absence of sialidase with added recombinant CTLA-4 (d) and (e) Anti-CD2 blocking antibody. (f) Staining of CD3ζ and F-actin to visualize the immune synapses formed by T cells and tumor cells by confocal microscopy. Two groups, with and without sialidase treatment of the tumor cells, were imaged. Scale bar =10 μm. (g) CD3 accumulation at the IS was calculated by dividing the mean fluorescence intensity (MFI) at the IS by the MFI of the rest of the membrane. (h) Relative IS contact area was calculated by dividing the area of the IS by the area of the rest of the T cell membrane. All analysis was done using ImageJ. Mean values show three independent experiments with standard error of the mean (SEM) as error bars. For statistical analysis, unpaired Student t test with Welch correction was applied (\**P* < 0.05, \*\**P* < 0.01, \*\*\**P* < 0.001 and \*\*\*\**P* < 0.0001).

Formation of BiTE-induced immunological synapse (IS) between target cells and T cells is the essential mode of action of BiTEs. We hypothesized that the removal of cell-surface sialosides may lead to stronger BiTE-induced IS formation between target tumor cells and T cells, and thus, promote better tumor cell killing. Accumulation of the TCR-CD3 complex and F-actin at the synapse is a hallmark of a stable and functional cytolytic IS in T cells. To test whether desialylation can enhance IS formation, we imaged the IS formed between T cells with sialidase-treated and non-treated SK-BR-3, a HER2-positive human breast cancer line, by staining F-actin and CD3ζ. The resulting immunofluorescence was imaged by confocal microscopy. As shown in Figure 2f, we observed visually larger BiTE-induced IS formation between T cells and desialylated SK-BR-3 cells. To assess the stability of the IS formed, we calculated the relative CD3 fluorescence intensity at the IS and the relative area of the IS. The IS formed by sialidase-treated tumor cells and T cells showed significantly stronger CD3 accumulation and larger IS contact area compared to IS formed by untreated tumor cells and T cells (Figure 2g and h). The same trend was observed for BiTE-induced IS formation between SKOV-3 cells and T cells, with stronger IS formed following sialidase treatment. (Figure S2). Furthermore, we also stained and incubated HER2 positive SK-BR-3 cells or CD19 positive NALM-6 cells with T cells under the treatment of 4D5 BiTE or CD19 BiTE with or without the desialylation. Cluster formation was then analyzed by FACS. It is observed that desialylation can significantly increase the cluster formation induced by BiTE between tumor cells and T cells (Figure S3a and b)

The interaction between CD2 and CD58 is known to play a critical role in the formation of a productive immunological synapse^22–24^. We found that the inhibition of this interaction with an anti-CD2 blocking antibody partially reversed the cytotoxicity enhancement from the sialidase addition, further suggesting that the desialylation triggers stronger target cell killing by facilitating a tighter interaction between target tumor cells and T cells (Figure 2e).

### HER2-targeting BiTE-sialidase fusion protein selectively desialylates HER2-positive cells

Having confirmed that sialidase treatment potentiates T cell-dependent tumor cell cytolysis induced by BiTE, we next sought to specifically direct sialidase to the tumor cell-T cell interface via BiTE conjugation. Confining sialidase activity to the target cells would potentiate tumor cell killing while limiting nonspecific desialylation of cells in the immune system. Importantly, sialyl-Lewis X, a sialylated tetrasaccharide, is essential for leukocyte tethering and rolling en route to sites of inflammation and tumor tissues. Nonspecific desialylation would destroy this glycan epitope on leukocytes, thereby hindering their tumor homing, and accordingly effective tumor control^25–27^. Toward this end, we constructed 4D5 BiTE–*B. infantis* sialidase fusion proteins in which sialidase was introduced onto either the N terminus (sialidase-4D5 BiTE) or the C terminus (4D5 BiTE-sialidase) of 4D5 BiTE, respectively (Figure 3a). To test whether the fusion protein can successfully remove sialic acids from the surface of tumor cells, SK-BR-3 (HER2+++) and SKOV-3, a human ovarian adenocarcinoma cell line (HER2+++) (Figure S4a, HER2+++ means high HER2 expression), were treated with Sialidase-4D5 BiTE or 4D5 BiTE-sialidase, respectively, followed by staining with the α-2,6-sialic acid-binding lectin SNA. Desialylation was evidenced by the decreased SNA binding as compared with the untreated controls and was seen in both SK-BR-3 and SKOV-3 cells when treated with either fusion protein (Figure 3b). To determine whether the sialidase fusion proteins can selectively desialylate HER2 positive cells in the presence of HER2 negative cells, we mixed SKOV-3 (HER2+++) and MDA-MB-468 (HER2-) cells, followed by the addition of 4D5 BiTE-sialidase. We observed that 4D5 BiTE-sialidase, at both 5 nM and 50 nM concentrations, selectively desialylates HER2 positive SKOV-3 cells while sparing the HER2 negative MDA-MB-468 cells, thus, confirming its selectivity for HER2-expressing cells (Figure 3c).

**Figure 3:**
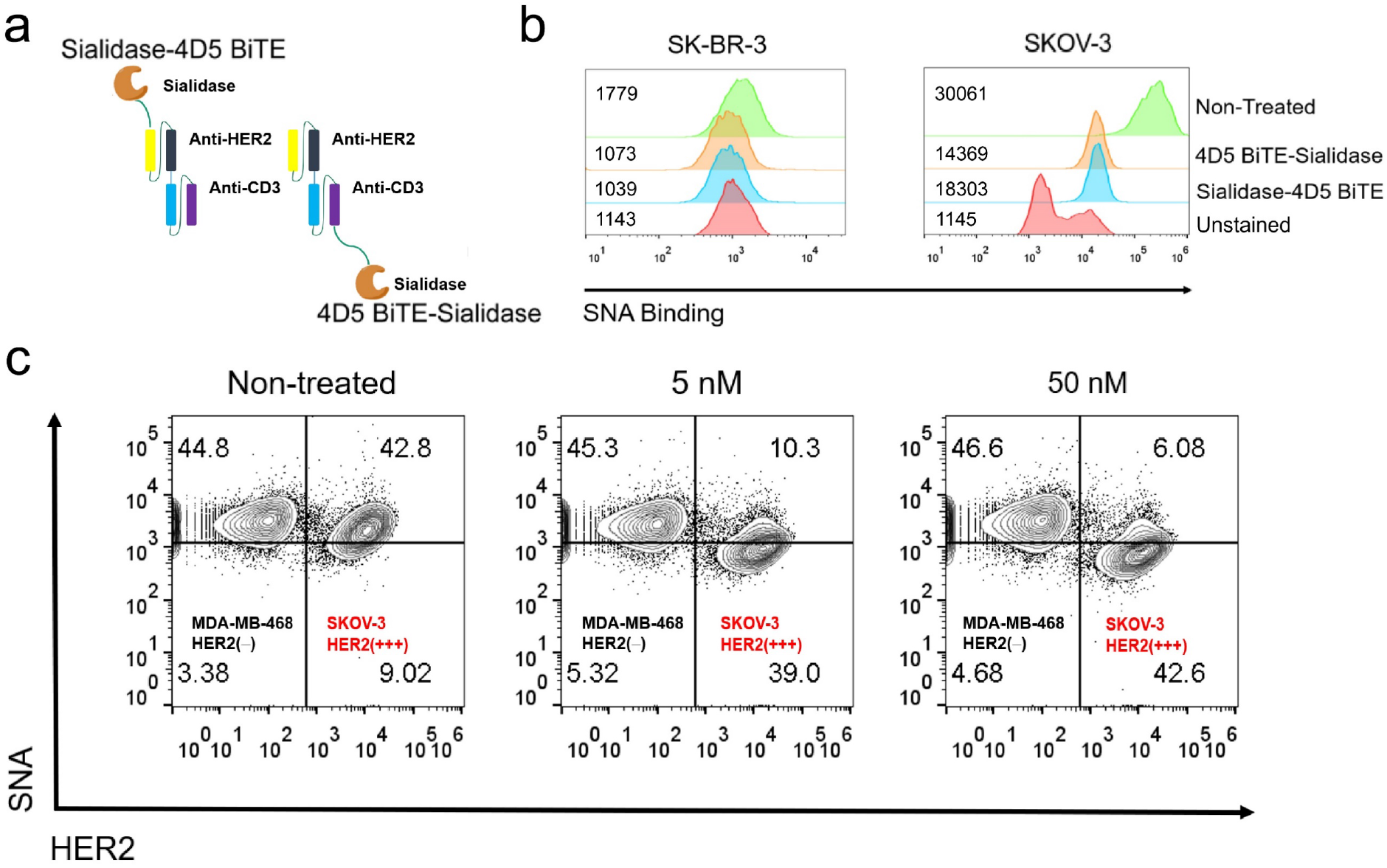
Construction of 4D5 BiTE-sialidase fusion proteins for selective desialylation of HER2 positive cells. (a) Two fusion proteins are constructed by conjugating sialidase to either the N or C terminus of the 4D5 BiTE. (b) Measuring sialic acid levels on the surface of SK-BR-3 and SKOV-3 cells after the treatment of the fusion proteins and staining with FITC-SNA. Mean fluorescent intensity (MFI) is displayed on the figure. (c) HER2 positive SKOV-3 cells and HER2 negative MDA-MB-468 cells were mixed and treated with 5 nM or 50 nM 4D5 BiTE-sialidase. The cell-surface sialylation level was measured by FITC-SNA staining and flow cytometry analysis.

### Anti-HER2 BiTE-sialidase triggers enhanced T cell-dependent *in vitro* cytotoxicity and T cell effector function than HER2 BiTE

We then compared the T cell-dependent cytotoxicity mediated by both fusion proteins to that of the original 4D5 BiTE. At the same concentration of 4 nM, both fusion proteins induced a higher level of T cell-dependent cytolysis of SK-BR-3 and SKOV-3 cells than 4D5 BiTE (Figure 4a and b). Specifically, in a dose-response assay, using SK-BR-3 and SKOV-3 cells as the target cells a tenfold and a threefold lower EC50, respective, were measured for 4D5 BiTE-sialidase than 4D5 BiTE (4D5 BiTE EC50 = ~200 pM) (Figure 4c and d). Consistent with these findings, 4D5 BiTE-sialidase induced the highest levels of T cell activation as measured by the expression of T cell activation markers CD25 and CD69 and the degranulation marker CD107a (Figure 4e to g). Also, the strongest release of cytokines, including IL-2, IFN-γ and TNF-α, was observed for 4D5 BiTE-sialidase-treated T cells. By contrast, sialidase-4D5 BiTE unexpectedly reduced the production of pro-inflammatory cytokines by T cells. (Figure 4h to j). A similar trend was seen for SKOV-3 cells, with 4D5 BiTE-sialidase inducing the strongest T cell activation (Figure S5). Due to its superior tumor cell killing efficacy, 4D5 BiTE-sialidase was chosen for further studies.

**Figure 4:**
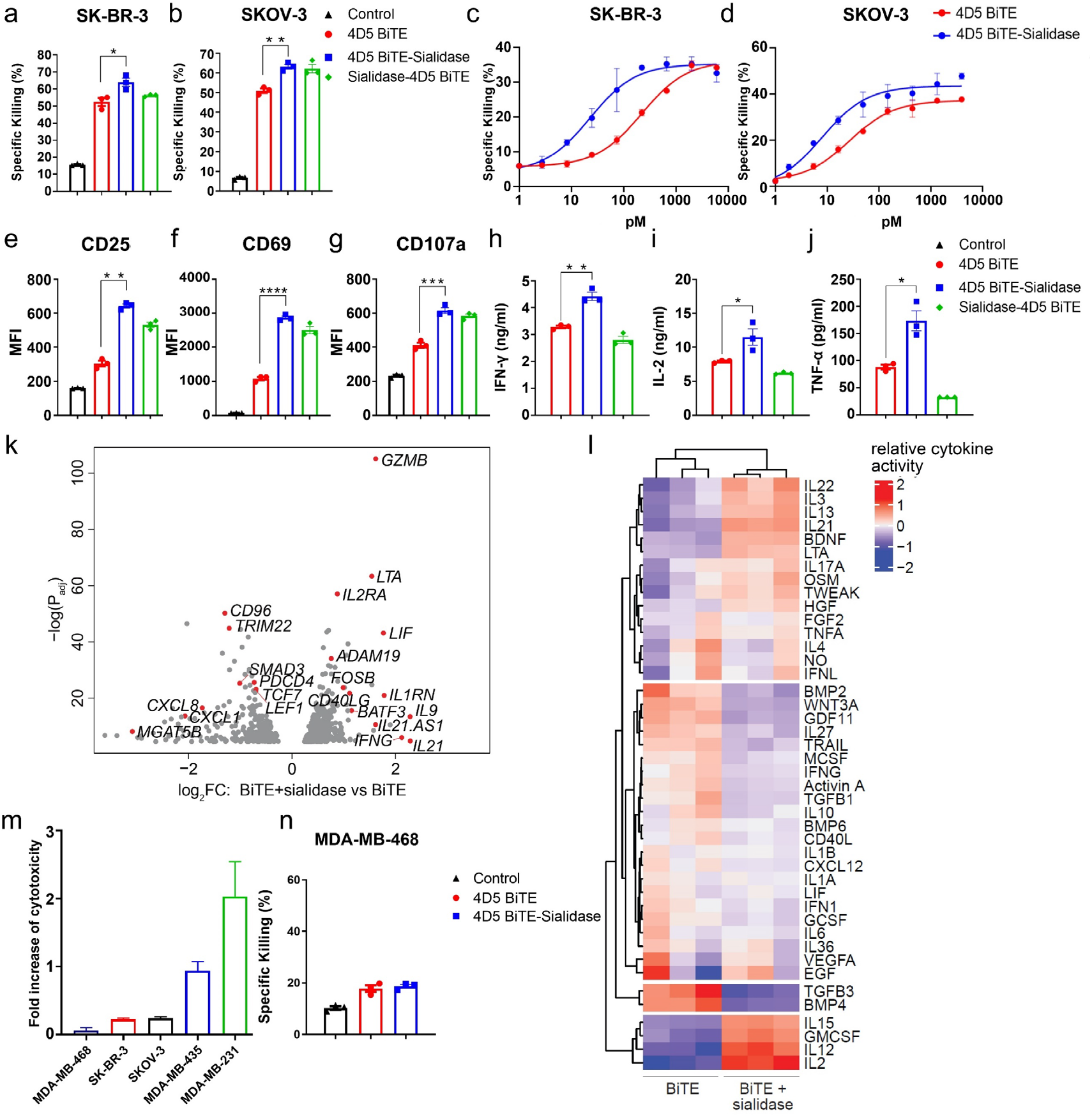
4D5 BiTE-sialidase fusion protein exhibits better activities than 4D5 BiTE alone for HER2 positive target cell killing and T cell activation. (a) and (b) The specific lysis of HER2 positive SK-BR-3 cells and SKOV-3 cells with 4 nM 4D5 BiTE or sialidase fusion proteins at the effector: target ratio of 5:1. (c) & (d) Dose dependent targeted killing using 4D5 BiTE or 4D5 BiTE-sialidase against SK-BR-3 and SKOV-3 cells. (e), (f) & (g) CD25, CD69 and CD107a expression level was measured in T cell populations in the presence of SK-BR-3 cells and 4 nM 4D5 BiTE or the fusion protein. (h), (i) & (j) IFN-γ, IL-2 and TNF-α release were measured for 4D5 BiTE- or fusion protein-induced T cell activation in the presence of SK-BR-3 cells. (k) Volcano plot of differentiated expressed genes between T cells treated with 4D5 BiTE or 4D5 BiTE-sialidase upon co-culturing with target MDA-MB-231 cells (genes with p value of fold changes < 0.01 were displayed). Relevant differentiated genes were highlighted in red. (l) The heatmap of cytokine activities ranked by Cytosig. (m) The cytotoxicity enhancements induced by 4D5 BiTE-sialidase as compared to 4D5 BiTE for cell lines with different HER2 expression levels. (n) The specific lysis of MDA-MB-468 cells under 4 nM 4D5 BiTE or 4D5 BiTE-sialidase at the effector: target ratio of 5:1. IC50 values were calculated from sigmoidal dose-response curve model using PRISM8. For statistical analysis, unpaired Student t test with Welch correction was applied (*P < 0.05, **P < 0.01, ***P < 0.001 and ****P < 0.0001).

The above studies showed that the 4D5 BiTE-sialidase engaged T cells are better activated versus those engaged by 4D5 BiTE. Therefore, it was of interest to determine whether the better T cell activation was originated from transcriptional alterations induced by BiTE treatment. To systematically characterize transcriptional changes in BiTE-molecule engaged T cells, we performed whole transcriptome RNA-sequencing (RNA-seq) analysis on either the 4D5 BiTE-sialidase or the 4D5 BiTE treated CD3^+^ T cells co-cultured with target MDA-MB-231 cells (Figure S6a). Volcano plot messenger RNA (mRNA) comparisons between the 4D5 BiTE-sialidase and the 4D5 BiTE treated T showed that 1191 transcripts were differentially expressed between these two groups (p < 3×10^−5^) (Figure 4k). The most highly expressed gene transcript in 4D5 BiTE-sialidase treated T cells encoded molecules crucial for T cell effector functions, including cytolytic enzymes and cytokines (*GZMB, LTA, LIF, IFNG*), cytokine receptors (*IL2RA*), and transcriptional regulators (*FOSB, BATF3*). Notably, gene transcripts associated with memory phenotypes, such as *LEF1* and *TCF7*, inhibitory receptors, such as *CD96* and *PDCD4*, and molecules involved in regulatory T cell generation, e.g., *SMAD3*, were largely downregulated. Gene-set-enrichment analysis (GSEA) highlighted multiple key pathways that are upregulated in the 4D5 BiTE-sialidase treated T cells, including those associated with cell cycle, transcriptional activity, and cell metabolism. Significantly, expression of transcripts involved in both oxidative phosphorylation and glycolysis was notably increased. By contrast, enrichment of downregulated genes pertained to pathways associated with Wnt-β catenin and TGF-β signaling (Figure S6b). Consistently, cytokine signaling (Cytosig) analysis revealed that pro-proliferation and inflammatory cytokines, IL-2, IL-12, IL-15, had the most clearly increased activity in the 4D5 BiTE-sialidase treated T cells, whereas the activity of the suppressive cytokine TGF-β3 was downregulated (Figure 4l)^28^. Together, compared with the T cells that were treated with 4D5 BiTE, the 4D5 BiTE-sialidase-treated T cells were in a more effector-differentiated state with higher oxidative phosphorylation, glycolysis activities and effector functions.

We further tested the BiTE-sialidase-mediated killing of cell lines with different cell-surface HER2 expression levels: MDA-MB-231 (+), MDA-MB-435 (+) and MDA-MB-468 (−) (Figure S5a). At 4 nM concentration, compared to 4D5 BiTE, 4D5 BiTE-sialidase strongly augmented the killing of cells with low levels of HER2 (HER2+), e.g., MDA-MB-231 and MDA-MB-435. Under this condition, stronger enhancements in killing were achieved than those measured for HER high (HER2+++) cells (SK-BR-3 and SKOV-3 cells) (94-203% *vs*. 22-24%) (Figure 4m and Figure S4c and d). These observations suggest that desialylation can increase the susceptibility of cells that would normally be relatively resistant to BiTE-mediated T cell killing. Significantly, 4D5 BiTE-sialidase did not trigger the killing of HER2 negative MDA-MB-468 cells or murine melanoma B16-F10 cells that express abundant sialoglycans, indicating exclusive specificity towards HER2 positive cells (Figure 4n and Figure S4b)

### BiTE-sialidase fusion proteins specific for CD19 and PSMA trigger enhanced *in vitro* cytotoxicity and T cell activation

To evaluate whether BiTE-sialidase fusion format can be applied to improve the efficacy of BiTE molecules targeting other tumor-associated antigens, we designed and constructed two additional BiTE-sialidase molecules. The first was based on the FDA-approved drug Blinatumomab that targets CD19, a cell surface marker on B cells and B cell malignancies. The second was derived from BiTE against prostate-specific membrane antigen (PSMA), a target for prostate cancer treatment. As shown in Figure 5a, compared to Blinatumomab (CD19 BiTE), the sialidase fusion counterpart exhibited much stronger cytotoxicity toward CD19-positive Raji cells with a fivefold lower EC50 (0.80 pM *vs*. 4.26 pM). Using the same concentration of 5 pM, CD19 BiTE-sialidase induced much higher T cell activation and degranulation than Blinatumomab (Figure 5b to e). Consistent with better T cell activation, CD19 BiTE-sialidase also triggered stronger cytokine release (Figure 5f to h). Likewise, much stronger killing of NALM-6, another CD19-positive cell line, was achieved by CD19 BiTE-sialidase (Figure S7). As what we observed for CD19 BiTE-sialidase, PSMA BiTE-sialidase also induced better killing of PC3 cells that were engineered to express high levels of PSMA and stronger T cell activation in comparison to PSMA BiTE (Figure S8a and b).

**Figure 5:**
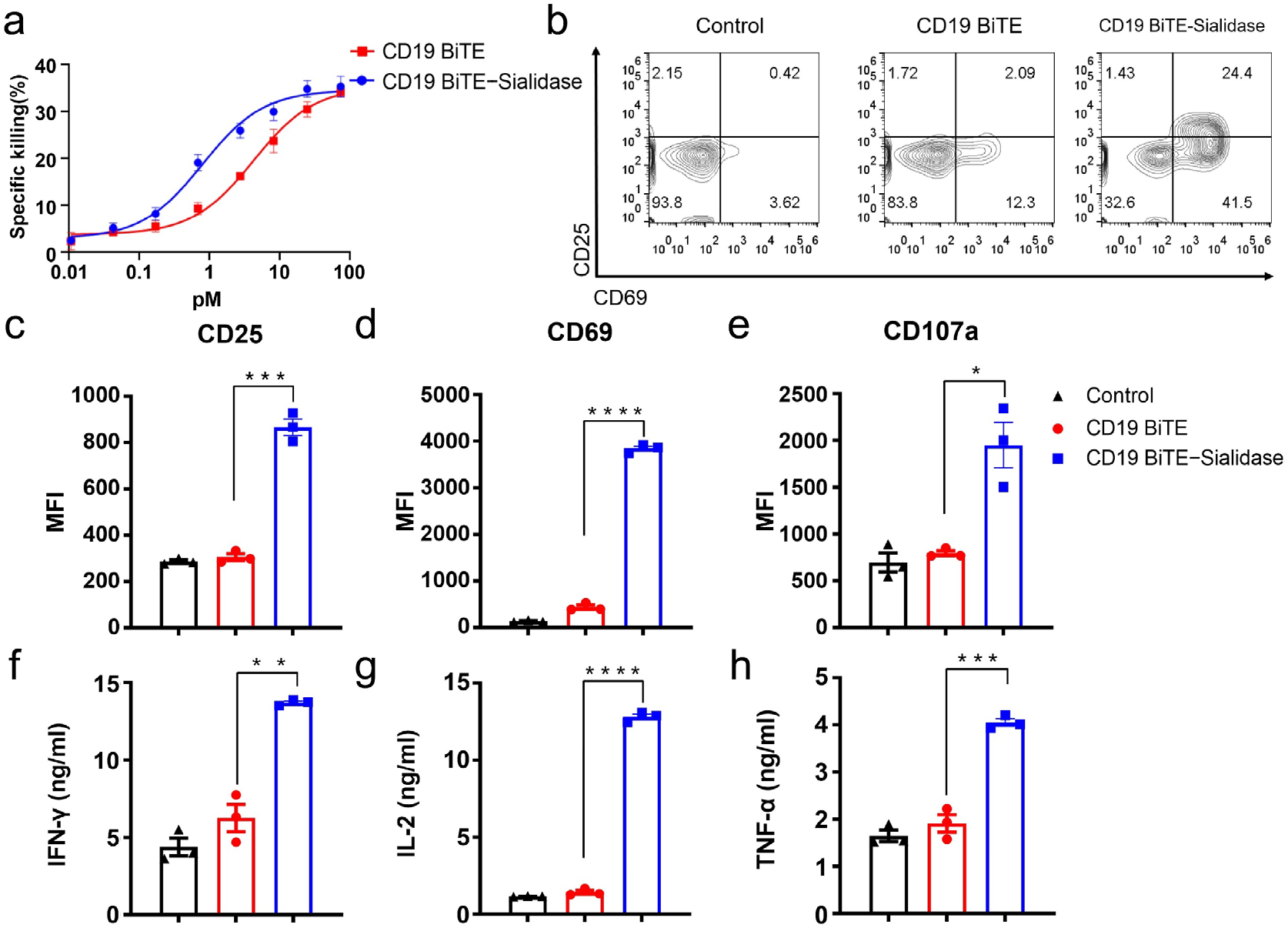
CD19 BiTE-sialidase exhibits superior activities than CD19 BiTE to induce *in vitro* tumor cell killing and T cell activation. (a) Dose-dependent killing of Raji cells induced by CD19 BiTE or CD19 BiTE-sialidase at the effector: target ratio of 10:1. (b), (c), (d) & (e) CD25, CD69, and CD107a expression levels were measured in T cell populations by flow cytometry analysis. (f), (g) & (h) IFN-γ, IL-2 and TNF-α release was measured for CD19 BiTE- or fusion protein-induced T cell activation in the presence of Raji cells. IC50 values were calculated from sigmoidal dose-response curve model using PRISM8. For statistical analysis, unpaired Student t test with Welch correction was applied (*P < 0.05, **P < 0.01, ***P < 0.001 and ****P < 0.0001).

### BiTE-sialidase enables better tumor control than BiTE in xenograft models in immune deficient mice

Having demonstrated the superiority of BiTE-sialidase fusion proteins versus the original BiTE molecules in terms of inducing T cell-dependent cytolysis of tumor cells *in vitro*, we then sought to determine if this enhanced efficacy could also be achieved *in vivo*. We chose a human tumor murine xenograft model using the NOD-*Prkdc*^em26Cd52^*IL2rg^em26Cd22^*/NjuCrl coisogenic (NCG) immunodeficient mouse to compare the antitumor immunity induced by 4D5 BiTE-sialidase and 4D5 BiTE constructs^29^. On day 0, NCG mice were injected subcutaneously (s.c.) with 2.5 million SK-BR-3-luc cells followed by intraperitoneal (i.p.) administration of 5 million hPBMCs. On day 7, these NCG mice were divided into three groups and then received an intravenous (i.v.) infusion of PBS, 4D5 BiTE, or 4D5 BiTE-sialidase, respectively (Figure 6a). After 5 hrs, blood was collected from each mouse to measure the serum IFN-γ level. The 4D5 BiTE-sialidase group was found to harbor the highest level of serum IFN-γ, whereas the 4D5 BiTE-treated group barely had any increased IFN-γ levels over the PBS control group (Figure 6b). The BiTE administration was continued twice per week until day 41. A second dose of 2 million hPBMCs per mouse was given on day 16. During this treatment course, tumor growth was monitored by longitudinal, noninvasive bioluminescence imaging. As shown in Figures 6c and 6d, the administration of 4D5 BiTE-sialidase significantly delayed tumor cell growth *in vivo* compared to the 4D5 BiTE treatment and PBS control. Remarkably, by the end of the treatment regimen, tumors in two mice receiving the 4D5 BiTE-sialidase treatment were completely eradicated (Figure 6e). We next investigated the *in vivo* efficacy of the CD19 BiTE-sialidase fusion protein using an orthotopic xenograft mouse model of leukemia. In this model, CD19^+^ NALM-6 cells (0.8 million) and hPBMCs (6 million) were injected intravenously (i.v.) into NCG mice on Day 0 (Figure 6f). The recipient mice were divided into four groups on day 3 and received i.v. infusion of PBS, 4D5 BiTE-sialidase, CD19 BiTE, or CD19 BiTE-sialidase, respectively. Significantly slower tumor progression was observed in the CD19 BiTE-sialidase treated group as compared to the CD19 BiTE treated group, demonstrating better *in vivo* antitumor effects of the sialidase fusion protein (Figure 6g and h). Notably, no apparent differences were detected between the PBS control group and the group that received non-CD19 targeting 4D5 BiTE-sialidase, indicating that the fusion protein triggered antitumor effects rely on the target engagement on tumor cells (Figure 6h).

**Figure 6:**
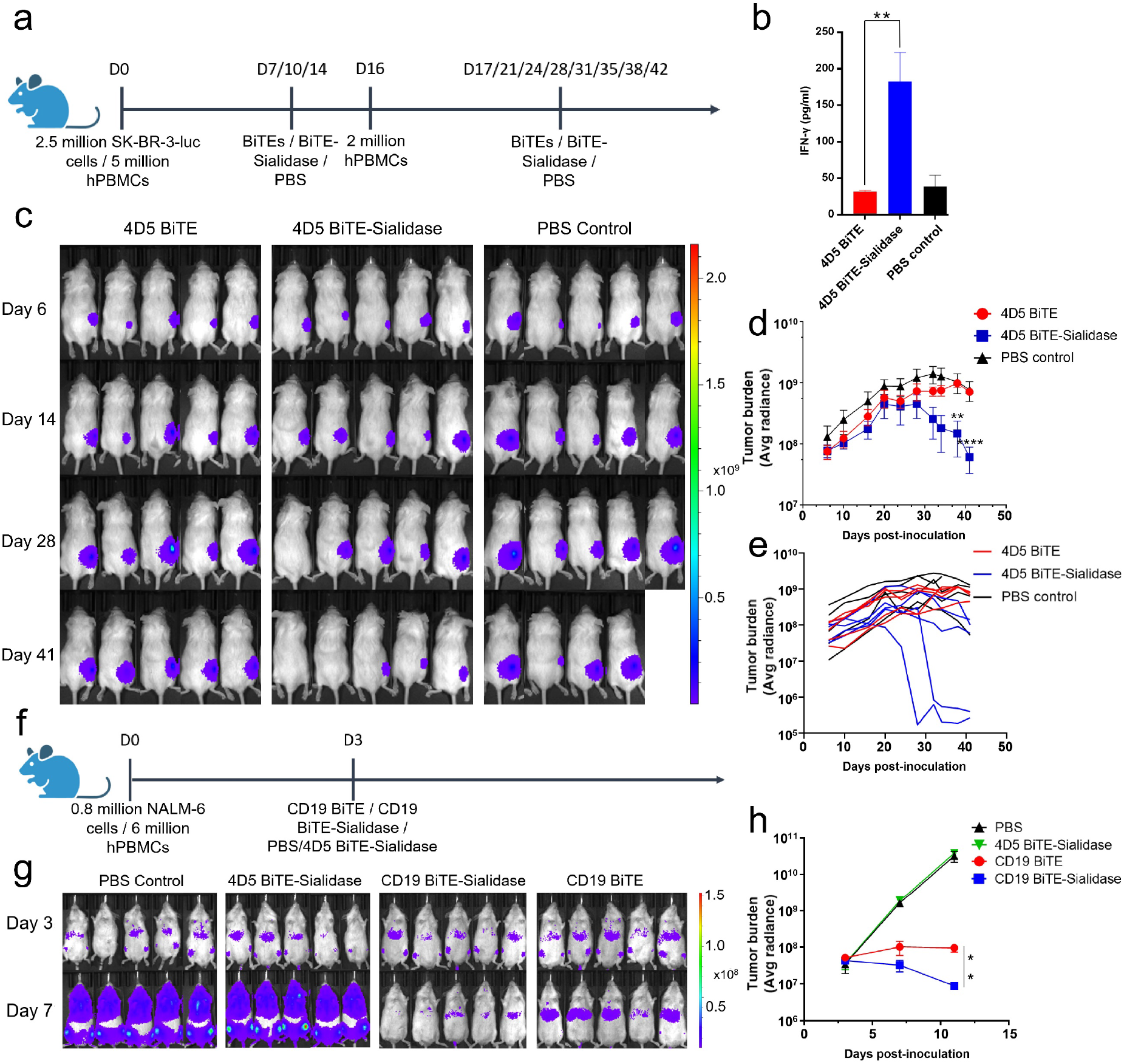
BiTE-sialidase fusion proteins exhibit better tumor control *in vivo* than BiTE. (a) Experimental timeline and treatment protocol for a HER2 positive SK-BR-3 breast cancer xenograft in NCG mice (n=5). (b) Serum IFN-γ release was measured 5 hours after the first drug treatment. (c) Bioluminescence was measured twice a week to visualize changes in tumor volume. On day 41, one mouse in the PBS control group died. (d) & (e) Bioluminescence was measured and calculated for each mouse as an indication of tumor burden. Tumor progression was followed by plotting change in the group average (d) and the individual (e) values over time. (f) Experimental timeline and treatment protocol for a xenograft NALM-6 model of acute lymphoblastic leukemia. (g) Bioluminescence measured on day 3 and day 7 were showed for different groups for comparing the tumor burden. (h) Tumor burden was measured and calculated by tracking the bioluminescence signals (n=5). One-way ANOVA and student t test were used to analyze the differences among groups (*P < 0.05, **P < 0.01, ***P < 0.001 and ****P < 0.0001).

### A BiTE-sialidase fusion protein showed therapeutic advantages over the parent BiTE in a syngeneic mouse model of melanoma

To further evaluate the efficacy of BiTE-sialidase fusion proteins in an immune-competent syngeneic mouse model, we constructed a murine CD3-engaging BiTE and the corresponding BiTE-sialidase from the ScFv fragments derived from anti-human EGFR antibody Cetuximab and anti-murine CD3ε clone 17A2. A mouse melanoma cell line, B16-EGFR5(B16-E5), with the expression of a chimeric mouse EGFR with six amino acid mutations to enable the binding of Cetuximab was chosen as the target cell. The fusion protein successfully induced desialylation of B16-E5 cells *in vitro* as confirmed by SNA staining (Figure 7a). To compare the anti-tumor activities of EGFR BiTE and EGFR BiTE-sialidase *in vivo*, we inoculated C57BL/6J mice with B16-E5 tumor cells (s.c) followed by intra-tumoral administration of EGFR BiTE or EGFR BiTE-sialidase. While both groups conferred therapeutic advantages over the PBS control group, the EGFR BiTE-sialidase treatment significantly delayed tumor growth as compared to the EGFR BiTE counterpart, in addition to offering notable survival benefits to the recipient mice (Figure 7b and c). Next, we investigated whether BiTE sialidase fusion protein conferred better tumor control by inducing changes in immune cell compositions in the tumor microenvironment. A single high dosage of EGFR BiTE or EGFR BiTE-sialidase was injected intratumorally on Day 11 post-tumor inoculation. Tumors and tumor-draining lymph nodes were harvested three days after the treatment (Figure 7d), at which point, the fusion protein-treated group had smaller tumor sizes compared to the BiTE treated group (Figure 7d). We found that in tumor-draining lymph nodes of both the EGFR BiTE and the EGFR BiTE-sialidase treated groups had significantly higher numbers of lymphocytes as compared with the PBS control group with the BiTE-sialidase treated group having the highest CD8^+^ T cell counts (Figure 7e and f). When analyzing tumor-infiltrating immune cells, compared with the EGFR BiTE treated groups, the BiTE-sialidase treated group had significantly higher frequencies of CD8^+^ T cells and NK cells (CD45.2^+^CD3^−^NK1.1^+^) and a reduced frequency of myeloid cells (CD45.2^+^CD11b^+^ NK1.1^−^) (Figure 7g to j). However, no apparent differences in CD4^+^ T cells and dendritic cells (CD45.2^+^CD11c^+^) were observed. We then analyzed CD8^+^ T cells in different groups and found that CD8^+^ T cells in the EGFR BiTE-sialidase treated group are skewed to a more effector-like phenotype (Figure S9a and b). Together, these results demonstrated that the BiTE-sialidase fusion protein facilitates the conversion of a myeloid-rich, T cell-poor tumor microenvironment that is immunosuppressive into a more immunopermissive one populated with NK and CD8^+^ T cells, which, in turn, leads to significantly improved tumor control.

**Figure 7:**
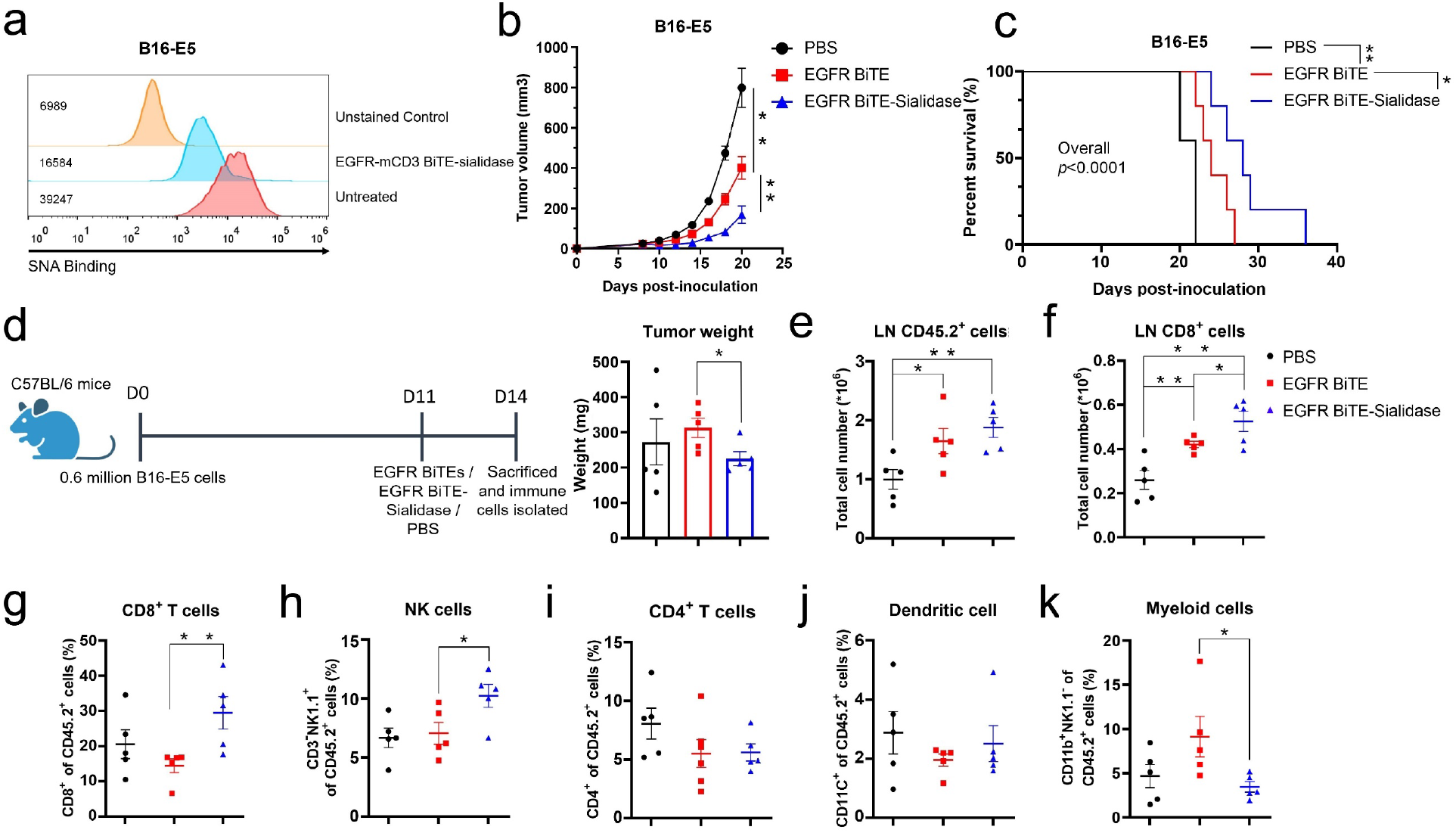
EGFR BiTE-sialidase shows better tumor control by altering tumor immune cell compositions in a syngeneic mouse model of melanoma. (a) Measuring sialic acid levels on the surface of B16-E5 cells after the treatment of EGFR BiTE-sialidase with sialic acid binding lectin SNA. Mean fluorescent intensity (MFI) is displayed on the figure (b) Tumor growth curve and percentage of survival (c) of B16-E5 mice model under intratumor treatment of PBS, EGFR BiTE and EGFR BiTE-sialidase. 0.6 million B16-E5 cells were inoculated in C57BL/6J mice (s.c.) and 0.5 μg BiTE or 0.93 μg BiTE-sialidase were given intratumorally on Day 8, 12 and 14, n=5 for each group. (d) Tumor infiltrated tumor cell profiling in B16-E5 mice model. (left) 0.6 million B16-E5 cells were inoculated in C57BL/6 J mice and 1.5 μg BiTE and 2.8 μg BiTE-sialidase were given intratumorally on Day 11 before being sacrificed on Day 14. (Right) Tumor weight at Day 14. (e) & (f) Total cell number count of CD45.2^+^ (e) and CD8^+^ (f) cells in the tumor lymph nodes isolated at Day 14. (g to k) the percentage of CD8 T cells (g), NK cells (h), CD4 T cells (i), CD11c dendritic cells (j) and CD11b myeloid cells (k) in the tumors of the different treatment groups (n=5). For statistical analysis, unpaired Student t test with Welch correction was applied. Kaplan-Meier with log-rank testing and Cox regression for survival analysis. (*P < 0.05, **P < 0.01, ***P < 0.001 and ****P < 0.0001)

## Discussion

The concept of cancer progression facilitated by hypersialylation was introduced more than five decades ago, but the idea of pursuing desialylation as a therapeutic strategy for cancer treatment has met with only mixed results since the earliest attempts in the 1970s^30–32^. Recently, the enthusiasm for this idea has been reinvigorated by the innovative work of Bertozzi and coworkers^33,34^. As demonstrated by Bertozzi, Laubli, *et al*., selective removal of sialylated Siglec-ligands in the tumor microenvironment using an antibody-sialidase conjugate enhanced anti-tumor immunity and suppressed tumor progression *in vivo* in several mouse tumor models^34,35^. Mechanistically, desialylation facilitated the conversion of tumor-associated macrophages with immunosuppressive phenotypes to their antitumoral counterparts^35^.

In the current study, we demonstrate that targeted desialylation is also beneficial to BiTE-based therapy, which so far has only had limited success in the treatment of solid tumors. BiTE-sialidase fusion proteins we developed here possess the capacity to trigger superior T cell-dependent cytotoxicity against target cancer cells both *in vitro* and *in vivo* in both solid tumor and hematologic cancer models. It is worth noting that the observed cytotoxicity enhancement of BiTEs resulting from desialylation seems to be independent of the inhibitory Siglec signaling pathways. Instead, desialylation elicits stronger immunological synapse formation between the engaged T cells and target cells. Indeed, several studies published recently revealed that bulky glycans on the surface of tumor cells result in suboptimal synapse formation induced by both CAR-T cells and BiTEs^36^. As a result of a better immunological synapse formation, T cells engaged by BiTE-sialidase fusion proteins exhibit increased activation, antigen sensitivity, and effector function.

Although we have evidence that the desialylation-induced cytotoxicity enhancement *in vitro* within a short window of 24 hours is independent of the Siglec signaling, recent studies showed that tumor-infiltrating T cells may express high levels of Siglec-9^16^. In addition, 48 hours following T-cell receptor stimulation, T cells upregulate Siglec-5 to counteract activation signals^37^. Therefore, desialylation by BiTE-sialidase fusion proteins may further facilitate long-term tumor cell killing by interrupting the inhibitory sialoglycan-Siglec interaction on T cells and other immune cells. Finally, the overexpression of sialosides on tumor cells is known to contribute to tumor metastasis^38–40^. There is a possibility that BiTEs-sialidase could suppress metastasis by causing targeted desialylation^41^. These hypotheses are currently under investigation in our lab.

To date, many efforts have been devoted to improving the efficacy of T cell-engaging therapies. One strategy is to include the CD28 costimulatory signal within the T cell-engaging process by either adding a CD28-engaging molecule or constructing trispecific molecules with both the CD3 and CD28 binding moieties^19–21^. Others have tried to incorporate immune checkpoint inhibitors (ICIs)^42^. BiTE-secreting CAR-T cells and BiTE-encoding oncolytic virus have also been developed in an effort to achieve better efficacy^43,44^. Distinct from these strategies, through the development of BiTE-sialidase fusion proteins we show that the efficacy of BiTEs can be significantly enhanced through targeted cell-surface glycocalyx editing. As illustrated in this work, the BiTE-sialidase fusion principle is applicable to BiTEs targeting various tumor-associated targets and therefore opens a new door for improving multiple types of T cell-engaging therapies. We anticipate that BiTE-sialidase fusion proteins will prove promising agents that can be combined with other anti-cancer modalities such as ICIs and adoptive cell transfer to achieve better tumor control.

## Methods

### Cell lines and cell culturing

SK-BR-3 cells, MCF7 cells, PC3 cells, Raji cells, SKOV-3 cells, MDA-MB-435 cells, MDA-MB-231 cells, MDA-MB-468 cells, NALM-6, NK92MI were obtained from ATCC and they were cultured as suggested. B16-E5 cells were kindly gifted from Yangxin Fu’s lab. Lentivirus transduction system was used to establish PSMA positive PC cells in which PSMA positive population was selected by FACs soriting. Expi293f cells were purchased from Thermo Fisher Scientific and cultured according to the protocol. For culturing of the isolated human PBMCs, AIM V™ Medium (Gibco™ 12055091) supplemented with 10% FBS was used. All cells were cultured in the incubator at 37°C supplemented with 5% CO2.

### General gene cloning procedures

The protein sequences of ScFv targeting human CD3, CD19, HER2 and PSMA were obtained from publicly available patents and the protein sequences were reverse-translated and codon-optimized to DNA sequences. All ScFv sequences were synthesized from IDT. The sequence of EGFR and murine CD3 binding ScFv were kindly provided by Yangxin Fu’s lab. The sequence of *B. infantis* sialidase was kindly gifted from Peng George Wang’s lab. For the molecular cloning process, the difference sequences were assembled using NEBuilder HiFi DNA Assembly (New England BioLabs, E2621). For BiTE molecules, two separate ScFv sequences were connected by a GGGGS linker. For the BiTE and sialidase fusion proteins, the sialidase sequence was conjugated to the BiTE sequence through a 2x GGGGS linker.

### Expression of BiTEs, *B. infantis* sialidase and BiTE-sialidase fusion proteins

All BiTEs, sialidase and BiTE-sialidase fusion proteins were fused with a 6x his tag at the C terminus for purification. For all BiTE and BiTE-sialidase fusion proteins, the expression was done in Expi293f cell system (Thermo Fisher Scientific). The transfection and handling of the cells were done according to the manufacturer’s protocol. *B. infantis* sialidase was expressed in BL21 *E. coli*. For purification, all proteins were purified using Ni-NTA (nickel-nitrilotriacetic acid) resin from QIAGEN. After the incubation of Expi293 media supernatant with the Ni-NTA resin, the Nickle charged resin was washed with PBS and 20 mM imidazole. Proteins were eluted with 250 mM imidazole and were concentrated and buffer-exchanged to PBS before use. The concentration of all proteins was determined by Qubit Protein Quantification Assay (Thermo Fisher Scientific, Q33211)

### Desialylation by *B. infantis* sialidase, 4D5 BiTE-sialidase fusion proteins and P-3Fax-Neu5Ac

For the removal of sialic acids by B. infantis sialidase or 4D5 BiTE-sialidase fusion proteins, 0.5 million cells were suspended in 100 mL DMEM without the serum. 1.5 mg sialidase or the fusion proteins were added in each sample and each sample was incubated at 37 °C for an hour. After the incubation, cells were washed twice by DPBS before they were used for killing experiments or staining. For the desialylation by inhibitor P-3Fax-Neu5Ac (R&D Systems, 117405-58-0), SK-BR-3 cells were cultured in T25 flask with the addition of 100 mM P-3Fax-Neu5Ac for three days.

### Desialylation detection from SNA staining

For the SNA staining, 0.5 million cells with or without desialylation were suspended in 100 μL HBSS buffer (Sigma-Aldrich, H6648) supplemented with 5 μM CaCl2 and MgCl2. SNA-FITC was added at 1:200 and DAPI was added at 1:2000 to each sample and the mixture was incubated on ice for 30 min before washing twice with HBSS buffer. Samples were then analyzed by FACS. Desialylation was analyzed in DAPI negative live cell populations using Flowjo.

### Human PBMC and T cell isolation

Human PBMCs were collected from blood samples of multiple healthy donors. Briefly, equal amount of DPBS with 2 mM EDTA was used to dilute the blood samples. Then, the mixture was carefully added to Ficoll (Ficoll® Paque Plus, GE Healthcare, 17-1440-02) for gradient separation. After centrifuge at 650g for 30 min with minimal acceleration and deceleration setting, the middle layer was collected and washed twice with DPBS supplemented with 2 mM EDTA. Further T cells isolation from human PBMCs was done with EasySep™ Human T Cell Isolation Kit (STEMCELL Technologies, 100-0695) according to the manufacturer’s protocol.

### Cell cytotoxicity measurement by lactate dehydrogenase (LDH) release

T cell cytotoxicity induced by BiTEs and BiTE-sialidase fusion proteins was measured by lactate dehydrogenase (LDH) release using CytoTox 96® Non-Radioactive Cytotoxicity Assay (Promega, G1780). 10000 tumor cells and 50000 hPBMCs per well in 100 μL media were exposed to different treatments and incubated in 96 well plates at 37 °C (unless different ratio was specified elsewhere). After 24 hours of coincubation, 50 μL of media supernatant from each well was transferred to a new flat bottom 96 well plate and LDH release was measured using the supplier’s protocol. Specific killing was calculated as suggested in the supplier’s protocol with background subtraction and total lysis comparison.

### Cytokine release and T cell surface activation marker measurement

For T cell cytokine release measurement, as with the cytotoxicity experiment, 10000 tumor cells and 50000 hPBMCs were co-incubated per well in 96 well plates with different treatments in 100 mL media at 37 °C for 24 hours. Then, 20 mL of supernatant from each well was diluted in 100 mL DPBS and used for IFN-γ, IL-2 and TNFα measurement. The ELISA measurement was done by ELISA MAX™ Sets. IFN-γ, IL-2 and TNFα kits (BioLegend) and the experiments were done according to the manufacturer’s protocol. The exact concentration was calculated from a standard curve. For the cell surface activation marker measurement, 80000 tumor cells and 400000 hPBMCs were co-incubated per well in 12 well plates with different treatments in 1 mL media at 37 °C for 24 hours. Following incubation, cells from each well were resuspended and stained with anti-CD3-PE, anti-CD69-FITC, anti-CD25-APC or anti-CD107a-pacific blue (All from biolegend and were added at 1:200) for 30 min at 4 °C. Cells were then washed twice with FACS buffer (PBS with 2.5% BSA) before being analyzed using flow cytometry. Data analysis and mean fluorescence intensity calculation were done by Flowjo. For the transcriptome analysis, 1.2 million hPBMCs and 0.1 million MDA-MB-231 cells were incubated together under the treatment of either 4 nM 4D5 BiTE or 4 nM 4D5 BiTE-sialidase (Three replicas for each condition).

After 48 hours of incubation, the mixture was stained with DAPI and CD3 to sort out the T cell population. mRNA of T cell from each population was extracted by The Arcturus PicoPure RNA Isolation Kit (Thermo fisher). The mRNA samples were sent out to Novogene for sequencing and initial analyzing.

### Flow cytometric analysis of Siglec-7 and Siglec-9 expression

Human PBMCs were collected from four healthy human donors. 0.5 million freshly isolated human PBMCs were suspended in 100 μL FACS buffer (PBS with 2.5% BSA) and each sample was stained with anti-CD3-PE. Each sample was also stained with either anti-Siglec-7-APC or anti-Siglec-9-APC (All from biolegend and were added at 1:200). After incubation for 30 min at 4 °C, cells were washed twice with FACS buffer before being analyzed using flow cytometry. Positive population percentage of both Siglec-7 and Siglec-9-stained samples was analyzed by Flowjo. For T cells activated by BiTEs, 80000 tumor cells and 400000 hPBMCs were coincubated per well in a 12 well plates with or without the BiTEs and sialidase treatment in 1 mL media at 37 °C for 24 hours. Following incubation, cells were resuspended and stained as described earlier for Siglec-7 and Siglec-9 expression analysis.

### Staining of human CD3ζ and actin for confocal imaging

Briefly, 0.4 million tumors cells were treated with 4 nM 4D5 BiTE or 4 nM 4D5 BiTE with 15 μg/ML sialidase in 100 μL DMEM without the serum for 1 hr at 37 °C. After the incubation, all the samples were washed twice using PBS before incubating with 0.4 million hPBMCs in 500 μl PBS for 30 min at 37 °C. Then all the cells were transferred in 1 ml PBS to the coverslips in 12 well plates and incubated at 37 °C for 30 min to let cells attach to the coverslip. 1 ml 4% PFA was added to each well and incubated with shaking for 20 min at room temperature (RT) for cell fixing, and then each well was washed twice with ice cold PBS. Washing took place at RT for 10 min with shaking. After fixation, 1 mL 0.1% PBS-Triton100 was added to each well for 10 min with shaking at RT to permeabilize the sample. PBST was used for washing for three times, each time with shaking at RT for 5 min. Next, 1 mL 2.5% FBS-PBST was used to block each sample for 50 min with shaking at RT. Then, anti-CD247(CD3ζ) antibody (Sigma-Aldrich, 12-35-22-00) was diluted in FACs buffer at 1:200 and anti-actin antibody (Novus Biologicals, NBP267113) was diluted in 1:500. 500 mL of each diluted antibody was added to samples and incubated for an hour at RT with shaking. PBST was used for washing for three times before anti-rabbit 488 (Invitrogen, 35553) and anti-mouse 594 ((Invitrogen, A-11005) secondary antibody was diluted and used for staining at RT for 30 min with shaking. Finally, samples were washed three times and each coverslip was transferred to a glass slide with mounting oil. Fingernail oil was used to seal the coverslip. Samples were analyzed on a Zeiss LSM880 with a 63x oil lens (NA 1.4). The relative mean fluorescent intensity (MFI) of CD3ζ accumulation and relative contact area of IS was calculated by imageJ.

### Cluster formation analysis

For the cluster formation experiments between SK-BR-3 cells and T cells. 0.5 million SK-BR-3 and 1 million hPMBCs were stained with CellTracker™ Green CMFDA (Thermo fisher) and PE anti-CD3 separately. After washing, they were incubated together under the treatment of the 50 nM 4D5 BiTE with or without the sialidase or 50 nM 4D5 BiTE-sialidase at 37 °C for 2 hours before the sample being analyzed by the FACS machine. The cluster experiment for the NALM-6 cells was of the same steps and settings except that NALM-6 GL cells carries GFP expression which doesn’t need CellTracker staining. All results were analyzed by Flowjo.

### RNA-sequencing analysis

Quality of raw sequencing reads was verified using FastQC (ref: Andrews, S. (2010). FastQC: A Quality Control Tool for High Throughput Sequence Data [Online]. Available online at: http://www.bioinformatics.babraham.ac.uk/projects/fastqc/). Reads were aligned to the genome and genic reads quantified using STAR version 2.7.0f^45^ and Ensembl version 101 GRCm38 genome and transcriptome annotations. Normalization, differential expression analysis and principal component analysis were performed using R package DESeq2 v1.35.0. Heatmaps were constructed using R package ComplexHeatmap v2.12.0. R version 4.2.1 was used. Cytokine target expression analysis was performed using the python implementation of CytoSig^28^. Gene set enrichment analysis was performed using GSEA^46^.

### Immunodeficient human tumor cell line xenograft mice model

All animal experiments were approved by the TSRI Animal Care and Use Committee. 15 NCG (6 weeks old male) mice (Charles Rivers Laboratories) were injected with 5 × 10^6^ human PBMCs (intraperitoneally) and 2.5 × 10^6^ SK-BR-3 cells (subcutaneously) on Day 0. On Day 6, mice were imaged by BLI and divided into groups based on similar tumor burden within each group. On Day 7, Three groups were intravenously (i.v) treated with PBS, 6 μg 4D5 BiTE, and 10 μg 4D5 BiTE-sialidase, respectively. Blood was collected from each mouse 5 hrs following BiTE administration and the serum IFN-γ level was measured using ELISA MAX™ (Biolengend). Drug treatment was continued twice a week, mouse received a second dose of 2 × 10^6^ human PBMCs (intraperitoneally) and each on day 16. Tumor burden was imaged multiple times throughout the whole study process. For the BLI imaging, 200 μL 15 g/L D-Luciferin, Potassium Salt (GoldBio) was injected intraperitoneally in each mouse and mice were imaged by IVIS imaging system (PerkinElmer) after 10 mins. For the NALM-6 model, 20 NCG (6 weeks old male) mice (Charles Rivers Laboratories) were injected with 6 × 10^6^ human PBMCs (i.v) and 0.8 × 10^6^ NALM-6 cells (i.v) on Day 0. On Day 3, all mice were imaged and divided into four groups. 1.5 μg CD19 BiTE, 2.8 μg CD19 BiTE-sialidase, 4D5 BiTE-sialidase and PBS were injected into different groups, respectively. Tumor size was measured by BLI like described earlier until the death of the PBS control group.

### B16-E5 syngeneic mice model

For the B16-E5 syngeneic mice model, 15 C57BL/6J mice (6 weeks old male) were injected with 0.6 × 10^6^ B16-E5 cells subcutaneously on day 0. On Day 8, tumor size was obtained by caliper measurement using the formula V = (W2 × L)/2 and mice were divided into different groups. Intratumor injection of 0.5 μg EGFR BiTE, 0.93 μg EGFR BiTE-sialidase and PBS were given to mice in different groups on Day 8, 12 and 14. Tumor size was recorded every two days until the mouse reached the endpoint of tumor size of 1000 mm^3^. For the tumor infiltrated lymphocytes profiling, 15 C57BL/6J mice (6 weeks old male) were also injected with 0.6 × 10^6^ B16-E5 cells subcutaneously on day 0. On Day 11, tumor size was measure and divided into three groups. 1.5 μg EGFR BiTE, 2.8 μg EGFR BiTE-sialidase and PBS were injected intratumorally into tumors in different groups. On Day 14, tumors were collected and tumor infiltrated lymphocytes from each tumor of different groups were stained with multiple markers for different populations within the CD45.2 lymphocytes for the profiling.

### Statistical analysis

Unless specified elsewhere, results are shown using GraphPad Prism version 8.0.0 with standard error of the mean (SEM) as error bars, each dot represents a biological replicate. P values were calculated using the built-in data analysis function of Microsoft excel or GraphPad Prism8.

## Author contributions

Z.Y., P.W. and R.A.L., conceived and designed research studies. Z.Y. and P.W. wrote the paper and analyzed the data. J.Z and J.R.T performed analysis on the transcriptome sequencing results. Z.Y., Y.H., G.G., C.W. and Y.S. performed all experiments described in the paper.

## Acknowledgements

This work was supported by the NIH (to P.W. R01AI154138 and to J.R.T and P.W. R01AI143884). J. Z. is a recipient of the Cancer Research Institute/Irvington postdoctoral fellowship.

## Reference

1 Goebeler, M.-E. & Bargou, R. C. T cell-engaging therapies—BiTEs and beyond. Nature Reviews Clinical Oncology 17, 418–434 (2020).

2 Przepiorka, D. et al. FDA approval: blinatumomab. Clinical Cancer Research 21, 4035–4039 (2015).

3 Arvedson, T. et al. Targeting Solid Tumors with Bispecific T Cell Engager Immune Therapy. Annual Review of Cancer Biology 6 (2021).

4 Singh, A., Dees, S. & Grewal, I. S. Overcoming the challenges associated with CD3+ T-cell redirection in cancer. British Journal of Cancer 124, 1037–1048 (2021).

5 Munn, D. H. & Bronte, V. Immune suppressive mechanisms in the tumor microenvironment. Current opinion in immunology 39, 1–6 (2016).

6 Hakomori, S. Glycosylation defining cancer malignancy: new wine in an old bottle. Proceedings of the National Academy of Sciences 99, 10231–10233 (2002).

7 Magalhães, A., Duarte, H. O. & Reis, C. A. Aberrant glycosylation in cancer: a novel molecular mechanism controlling metastasis. Cancer cell 31, 733–735 (2017).

8 Pinho, S. S. & Reis, C. A. Glycosylation in cancer: mechanisms and clinical implications. Nature Reviews Cancer 15, 540–555 (2015).

9 Dube, D. H. & Bertozzi, C. R. Glycans in cancer and inflammation—potential for therapeutics and diagnostics. Nature reviews Drug discovery 4, 477–488 (2005).

10 Pearce, O. M. & Läubli, H. Sialic acids in cancer biology and immunity. Glycobiology 26, 111–128 (2016).

11 Rodrigues, E. & Macauley, M. S. Hypersialylation in cancer: modulation of inflammation and therapeutic opportunities. Cancers 10, 207 (2018).

12 Murugesan, G., Weigle, B. & Crocker, P. R. Siglec and anti-Siglec therapies. Current opinion in chemical biology 62, 34–42 (2021).

13 Smith, B. A. & Bertozzi, C. R. The clinical impact of glycobiology: targeting selectins, Siglecs and mammalian glycans. Nature Reviews Drug Discovery 20, 217–243 (2021).

14 Edgar, L. J. et al. Sialic acid ligands of CD28 suppress costimulation of T cells. ACS central science 7, 1508–1515 (2021).

15 Büll, C. et al. Sialic acid blockade suppresses tumor growth by enhancing T-cell–mediated tumor immunity. Cancer research 78, 3574–3588 (2018).

16 Stanczak, M. A. et al. Self-associated molecular patterns mediate cancer immune evasion by engaging Siglecs on T cells. The Journal of clinical investigation 128, 4912–4923 (2018).

17 Macauley, M. S., Crocker, P. R. & Paulson, J. C. Siglec-mediated regulation of immune cell function in disease. Nature Reviews Immunology 14, 653–666 (2014).

18 Hudak, J. E., Canham, S. M. & Bertozzi, C. R. Glycocalyx engineering reveals a Siglec-based mechanism for NK cell immunoevasion. Nature chemical biology 10, 69–75 (2014).

19 Wu, L. et al. Trispecific antibodies enhance the therapeutic efficacy of tumor-directed T cells through T cell receptor co-stimulation. Nature Cancer 1, 86–98 (2020).

20 Correnti, C. E. et al. Simultaneous multiple interaction T-cell engaging (SMITE) bispecific antibodies overcome bispecific T-cell engager (BiTE) resistance via CD28 co-stimulation. Leukemia 32, 1239–1243 (2018).

21 Skokos, D. et al. A class of costimulatory CD28-bispecific antibodies that enhance the antitumor activity of CD3-bispecific antibodies. Science translational medicine 12, eaaw7888 (2020).

22 Leitner, J., Herndler-Brandstetter, D., Zlabinger, G. J., Grubeck-Loebenstein, B. & Steinberger, P. CD58/CD2 is the primary costimulatory pathway in human CD28− CD8+ T cells. The Journal of Immunology 195, 477–487 (2015).

23 Demetriou, P. et al. A dynamic CD2-rich compartment at the outer edge of the immunological synapse boosts and integrates signals. Nature immunology 21, 1232–1243 (2020).

24 Shen, Y. et al. Cancer cell-intrinsic resistance to BiTE therapy is mediated by loss of CD58 costimulation and modulation of the extrinsic apoptotic pathway. Journal for immunotherapy of cancer 10 (2022).

25 Ley, K. & Kansas, G. S. Selectins in T-cell recruitment to non-lymphoid tissues and sites of inflammation. Nature Reviews Immunology 4, 325–336 (2004).

26 Sackstein, R., Schatton, T. & Barthel, S. R. T-lymphocyte homing: an underappreciated yet critical hurdle for successful cancer immunotherapy. Laboratory investigation; a journal of technical methods and pathology 97, 669–697, doi:10.1038/labinvest.2017.25 (2017).

27 Sackstein, R. Glycosyltransferase-programmed stereosubstitution (GPS) to create HCELL: engineering a roadmap for cell migration. Immunological reviews 230, 51–74, doi:10.1111/j.1600-065X.2009.00792.x (2009).

28 Jiang, P. et al. Systematic investigation of cytokine signaling activity at the tissue and single-cell levels. Nature methods 18, 1181–1191 (2021).

29 Eberle, C. et al. in *JOURNAL FOR IMMUNOTHERAPY OF CANCER*. (BMC CAMPUS, 4 CRINAN ST, LONDON N1 9XW, ENGLAND).

30 Bekesi, J. G., Arneault, G. S., Walter, L. & Holland, J. F. Immunogenicity of leukemia L1210 cells after neuraminidase treatment. Journal of the National Cancer Institute 49, 107–118 (1972).

31 Bekesi, J. G., Roboz, J. P. & Holland, J. F. Therapeutic effectiveness of neuraminidase-treated tumor cells as an immunogen in man and experimental animals with leukemia. Annals of the New York Academy of Sciences 277, 313–331 (1976).

32 Sedlacek, H., Hagmayer, G. & Seiler, F. Tumor therapy of neoplastic diseases with tumor cells and neuraminidase. Cancer Immunology, Immunotherapy 23, 192–199 (1986).

33 Xiao, H., Woods, E. C., Vukojicic, P. & Bertozzi, C. R. Precision glycocalyx editing as a strategy for cancer immunotherapy. Proceedings of the National Academy of Sciences 113, 10304–10309 (2016).

34 Gray, M. A. et al. Targeted glycan degradation potentiates the anticancer immune response in vivo. Nature Chemical Biology 16, 1376–1384, doi:10.1038/s41589-020-0622-x (2020).

35 Stanczak, M. A. et al. Targeting cancer glycosylation repolarizes tumor-associated macrophages allowing effective immune checkpoint blockade. BioRxiv, doi:doi.org/10.1101/2021.04.11.439323 (2021).

36 Greco, B. et al. Disrupting N-glycan expression on tumor cells boosts chimeric antigen receptor T cell efficacy against solid malignancies. Sci Transl Med 14, eabg3072, doi:10.1126/scitranslmed.abg3072 (2022).

37 Vuchkovska, A. et al. Siglec-5 is an inhibitory immune checkpoint molecule for human T cells. Immunology, doi:10.1111/imm.13470 (2022).

38 Dobie, C. & Skropeta, D. Insights into the role of sialylation in cancer progression and metastasis. British Journal of Cancer 124, 76–90 (2021).

39 Yogeeswaran, G. & Salk, P. Metastatic potential is positively correlated with cell surface sialylation of cultured murine tumor cell lines. Science 212, 1514–1516 (1981).

40 Bresalier, R. S. et al. Enhanced sialylation of mucin-associated carbohydrate structures in human colon cancer metastasis. Gastroenterology 110, 1354–1367 (1996).

41 Bull, C. et al. Targeted delivery of a sialic acid-blocking glycomimetic to cancer cells inhibits metastatic spread. ACS nano 9, 733–745 (2015).

42 Herrmann, M. et al. Bifunctional PD-1× αCD3× αCD33 fusion protein reverses adaptive immune escape in acute myeloid leukemia. Blood, The Journal of the American Society of Hematology 132, 2484–2494 (2018).

43 Choi, B. D. et al. CAR-T cells secreting BiTEs circumvent antigen escape without detectable toxicity. Nature biotechnology 37, 1049–1058 (2019).

44 Heidbuechel, J. P. & Engeland, C. E. Oncolytic viruses encoding bispecific T cell engagers: a blueprint for emerging immunovirotherapies. Journal of hematology & oncology 14, 1–24 (2021).

45 Dobin, A. et al. STAR: ultrafast universal RNA-seq aligner. Bioinformatics 29, 15–21 (2013).

46 Subramanian, A. et al. Gene set enrichment analysis: a knowledge-based approach for interpreting genome-wide expression profiles. Proceedings of the National Academy of Sciences 102, 15545–15550 (2005).

