## Supplemental figures for "Enhancing the anti-tumor efficacy of Bispecific T cell engagers via cell surface glycocalyx editing"

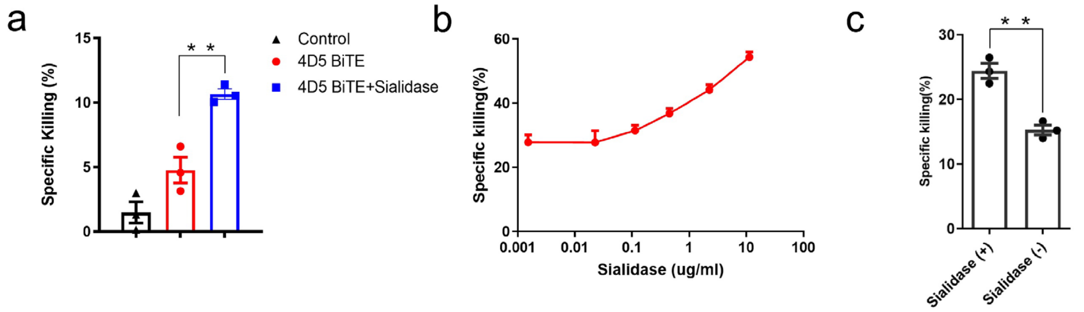


Figure S1: Desialylation enhances BiTE-induced T cell cytotoxicity. (a) Testing BiTE triggered killing against SK-BR-3 cells (E:T = 1:1) with or without the sialidase treatment with hPBMCs. (b) Different concentrations of sialidase were added with 4D5 BiTEs to test killing of SK-BR-3 cells by hPBMCs. (c) Isolated pure T cells were used to test killing of MCF-7 cells induced by BiTEs with or without the addition of 15 ug/ml sialidase. Unpaired Student t test with Welch correction was applied to analyze the data (*P < 0.05, **P < 0.01, ***P < 0.001 and ****P < 0.0001, ns = not important).

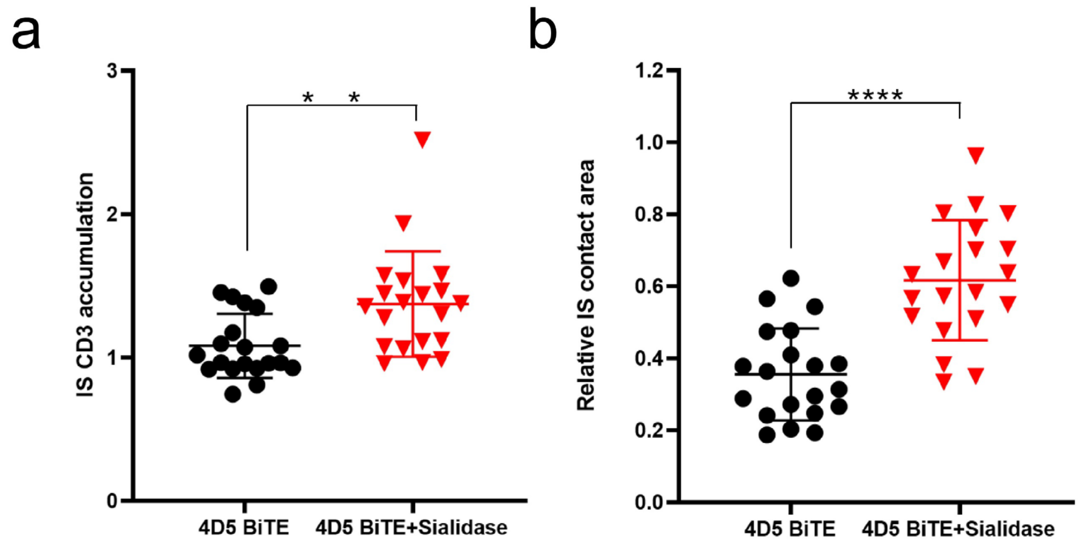


Figure S2: 4D5 BiTE induces stronger immunological synapse formation between desialylated SKOV-3 cells and T cells. (a) CD3 accumulation at the IS was calculated by dividing the mean fluorescence intensity (MFI) at the IS by the MFI of the rest of the membrane. (b) Relative IS contact area was calculated by dividing the area of the IS by the area of the rest of the T cell membrane. All analysis was done using ImageJ. For statistical analysis, unpaired Student t test with Welch correction was applied (*$P$ < 0.05, **$P$ < 0.01, ***$P$ < 0.001 and ****$P$ < 0.0001).

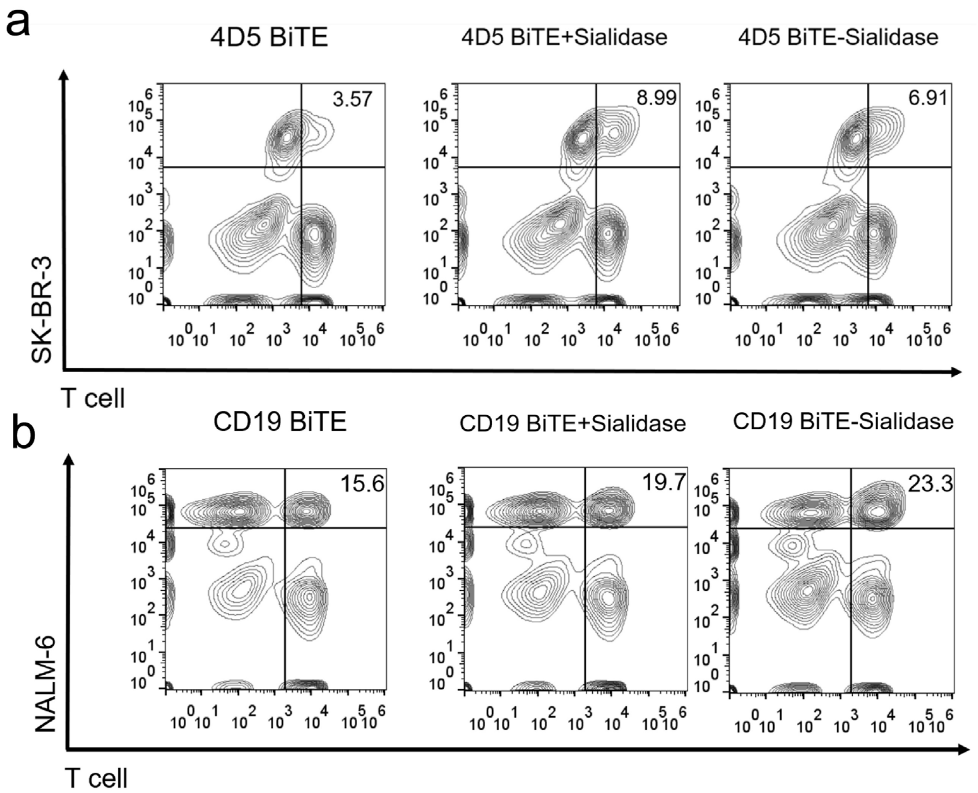


Figure S3: Stronger cluster formation is induced by BiTE between tumor cells and T cell with desialylation. Cluster formation measured between SK-BR-3 cells (a), or NALM-6 cells (b) and T cells was measured by FACS under corresponding BiTE treatment with or without the desialylation.

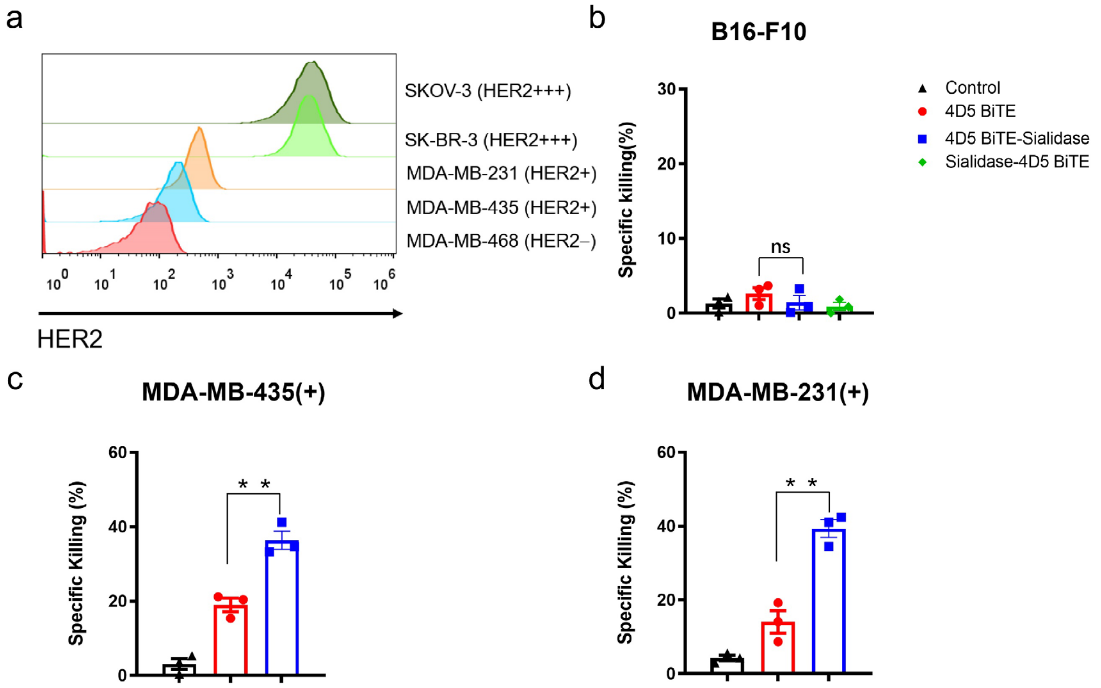


Figure S4: In vitro cytotoxicity of 4D5 BiTE-sialidase against tumor cell lines with different HER2 expression levels. (a) Measuring of HER2 expression levels on the surface of different human cancer cell lines by flow cytometry. (b), (c) and (d) The specific lysis of B16-F10, MDA-MB-435 and -231 cells under 4 nM 4D5 BiTE or the sialidase fusion protein at the effector: target ratio of 5:1. For statistical analysis, unpaired Student t test with Welch correction was applied (*P < 0.05, **P < 0.01, ***P < 0.001 and ****P < 0.0001).

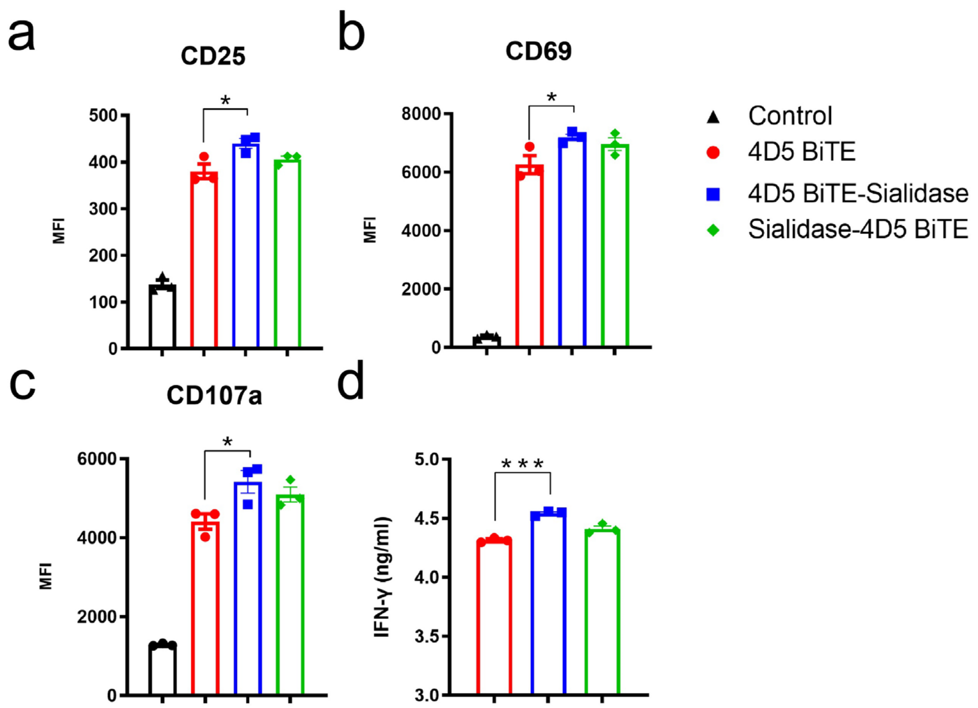


Figure S5: Characterization of T cell activation and cytokine release triggered by sialidase-4D5 BiTE fusion protein in the presence of target SKOV-3 cells. (a), (b) & (c) CD25, CD69 and CD107a expression level was measured in T cell populations in the presence of SKOV-3 cells and 4nM 4D5 BiTE or the Sialidase fusion proteins. (d) IFN-γ release was measured for 4D5 BiTE- or fusion protein-induced T cell activation in the presence of SKOV-3cells. For statistical analysis, unpaired Student t test with Welch correction was applied (*$P < 0.05$, **$P < 0.01$, ***$P < 0.001$ and ****$P < 0.0001$).

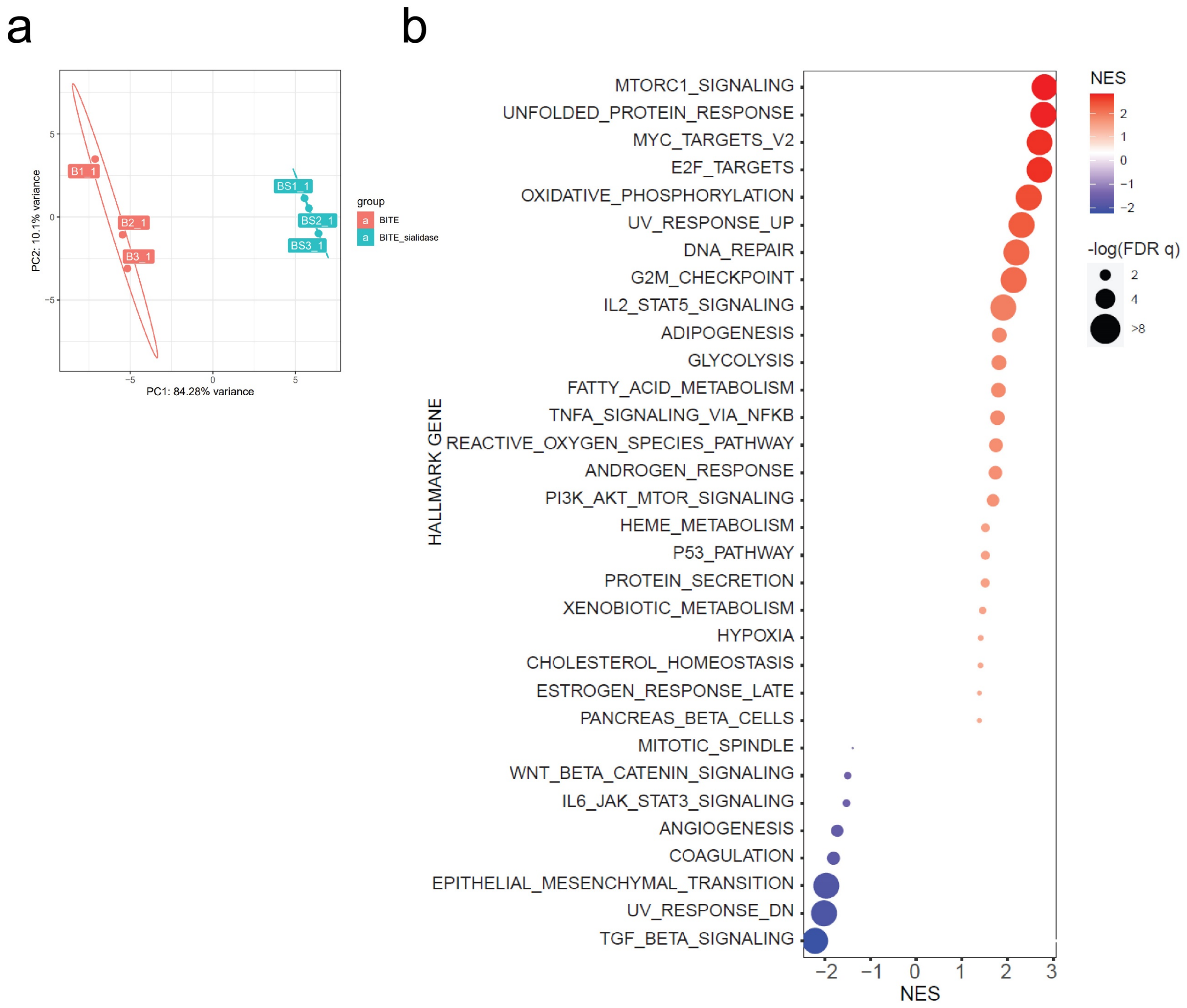


Figure S6: Global transcriptome analysis of T cell treated with 4D5 BiTE or 4D5 BiTE-sialidase upon co-culturing with target MDA-MB-231 cells. (a) Principal component plot with sample labels added. Top 1000 genes by variance used to construct principal components. (b) Gene set enrichment analysis of T cell group treated with 4D5 BiTE-sialidase versus 4D5 BiTE.

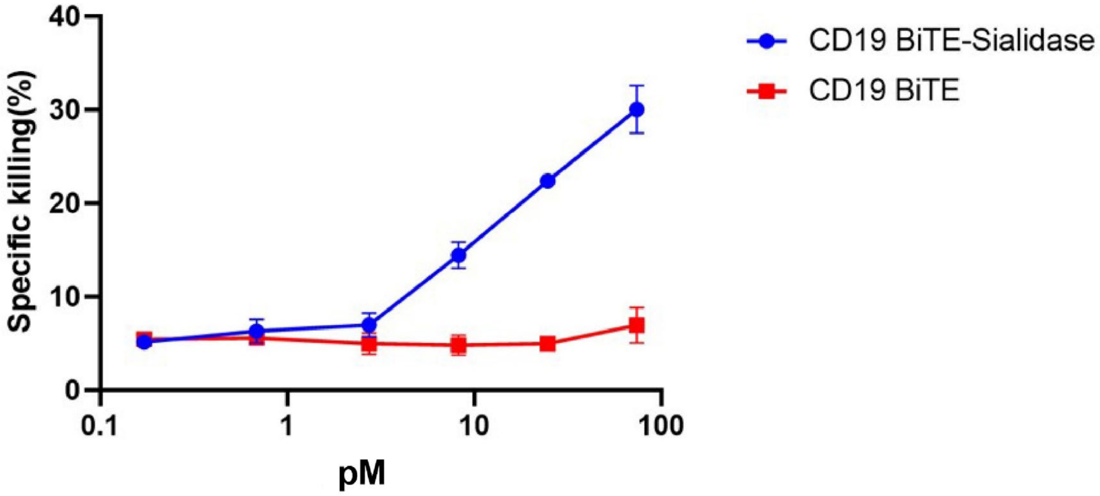


Figure S7: Testing the killing of NALM-6 cells under different concentrations of CD19 BiTE and CD19 BiTE-Sialidase at the effector: target ratio of 5:1.

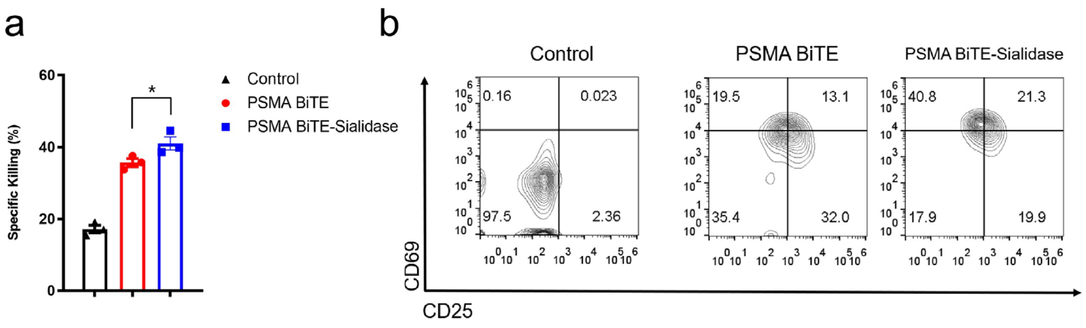


Figure S8: Measurement of the PSMA BiTE and BiTE-Sialidase mediated *in vitro* tumor cell killing and T cell activation. (a) The specific lysis of PSMA positive PC3 cells with 4nM PSMA BiTE or PSMA BiTE-Sialidase at the effector: target ratio of 5:1. (b) CD25 and CD69 expression levels were measured in the T cell population against PSMA positive PC3 cells with 4 nM PSMA BiTE or the Sialidase fusion protein. For statistical analysis, unpaired Student t test with Welch correction was applied (*$P < 0.05$, **$P < 0.01$, ***$P < 0.001$ and ****$P < 0.0001$).

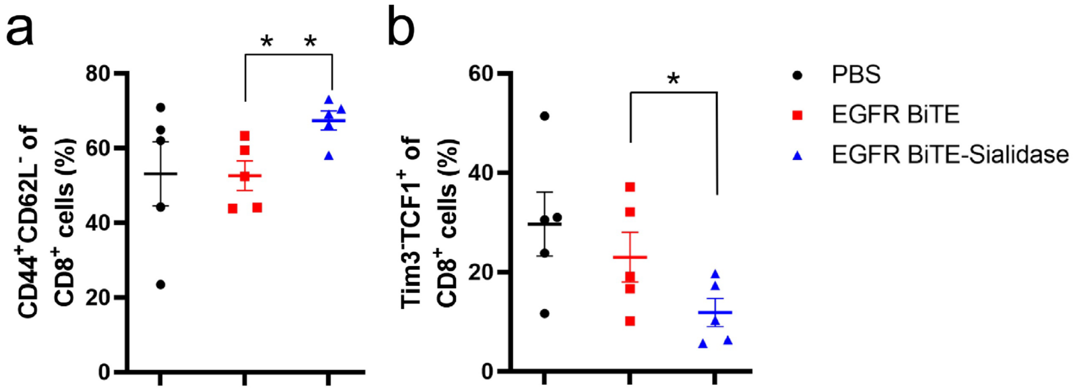


Figure S9: Profiling of the CD8 T cell in tumors treated with PBS, EGFR BiTE or EGFR BiTE-sialidase. CD44, CD61L (a), Tim3 and TCF1 (b) were stained among CD8 T cell group from collected tumors of different groups to profile CD8 T cell functionality. For statistical analysis, unpaired Student t test with Welch correction was applied (*P < 0.05, **P < 0.01, ***P < 0.001 and ****P < 0.0001).

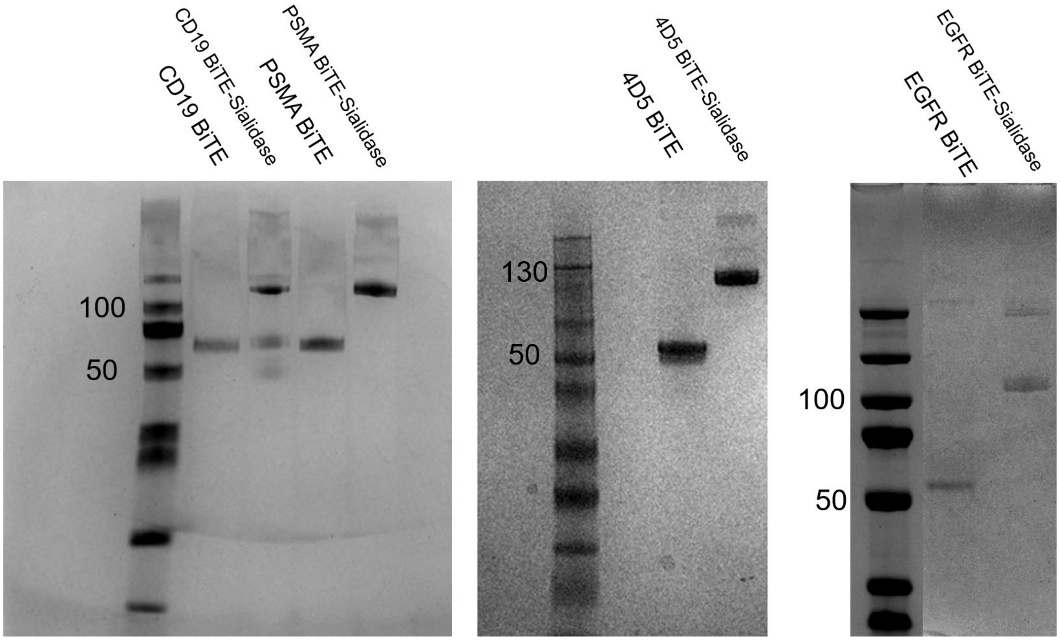


Figure S10：SDS-PAGE analysis of BiTE and BiTE-sialidase fusion proteins.
